# RADIX: a deep learning framework that maps root barriers across species and reveals genetic and environmental contributions

**DOI:** 10.64898/2026.08.07.743584

**Authors:** Yifei Gu, Stefan Sanow, Tamera Taylor, Kevin W. Morimoto, Adele Nemer, Dustin J. Hadley, Syed Adeel Zafar, Lucas DeMello, Yubei Chen, Hannah Knab, Axel Busch Castro, Varun Kumaravelu, Julia Bailey-Serres, Randy Carney, Siobhán M. Brady

## Abstract

Root anatomical barriers, including the suberized and lignified walls of the endodermis and exodermis, and cortical aerenchyma, regulate water and nutrient transport, gas exchange, and rhizosphere interaction. Their adaptive function places them as an important target for breeding environmentally resilient plant species. Quantifying these structures at high resolution is a manual bottleneck that limits experimental scale. We present RADIX (Root Anatomy Deep-learning Image segmentation across species and platforms), a framework that adapts a large self-supervised vision-transformer foundation encoder (DINOv3), pre-trained on billions of natural images, to root anatomy by fine-tuning its encoder with a dense-prediction-transformer decoder. Transferring these general-purpose vision encoders to a specialized biological domain with a high-quality annotated dataset is what allows RADIX to generalize across species and imaging platforms. We train and evaluate it on the first expert-annotated benchmark of root anatomical structures at scale, comprising 1,695 high-quality fluorescence images spanning 17 monocot and dicot species, six anatomical structures, and three imaging platforms. RADIX segments all six structures at inter-annotator-level accuracy and generalizes to unseen species, genotypes, growth conditions, and an imaging platform from an independent laboratory. A single unified model surpasses monocot- and dicot-specialist models without sacrificing in-group accuracy. Predicted masks yield aerenchyma and suberin/lignin measurements matching expert annotation at ∼1.2 s per image with a single GPU, reducing weeks of manual analysis to minutes. Applying RADIX across genotypes, microbial treatments, and growth systems, we show that these cell type features form a coordinated, multidimensional, and context-dependent system shaped by genetic and environmental factors.

## Introduction

Roots are the hidden half of plants, and their position below ground has precluded facile understanding of their form and function. High-throughput imaging and automated quantification of root system architecture have advanced rapidly, but quantification of root anatomy at cellular resolution remains comparatively difficult. Root cell types are arranged in radially symmetric, concentric cylinders surrounding the central vasculature. From the outside in, the epidermis is followed by the exodermis (when present), the cortex, and the endodermis, while the inner stele comprises the pericycle together with xylem, phloem, and procambial tissue. This radial patterning is broadly conserved, yet the number of layers and their morphology vary across plant lineages. *Arabidopsis thaliana*, the predominant model species, has a single cortical layer, whereas most species have multiple, often specialized, cortical layers. Functional specialization is frequently conferred by deposition of complex cell wall barriers of lignin, suberin, and other biopolymers, which control water and mineral ion uptake and exclude pathogens, while their localized absence can permit symbiont entry^1–7^. The exodermis, the outermost cortical layer, can be lignified and/or suberized, and inner cortical layers can accumulate morphologically diverse lignin deposition^8–10^. The endodermis contains a conserved, lignified Casparian strip and can additionally become suberized as well as a target of the microbiome^2,3,11^. Many monocot roots also form aerenchyma, gas-filled lacunae produced by cell-wall loosening and programmed cell death, that lower the metabolic cost of soil exploration and sustain internal oxygen supply under hypoxic or flooded conditions^12,13^. These features are dynamically remodeled in response to abiotic and biotic stress, and their increased deposition or abundance is often associated with stress tolerance^14^. A central question remains: what is the full diversity of these cell types and their differentiation, and how does it enable adaptive responses to the environment?

Resolving these cell types, cell wall barriers, and aerenchyma requires thin sectioning and cellular resolution imaging. Laser ablation tomography resolves cortical cell-wall features effectively and has been used to define the genetic architecture, function, and environmental significance of these traits in maize, wheat, and their wild relatives, but the instrumentation is not widely accessible^9,13,15^. Transverse sections are more readily obtained with a vibratome from fresh or agar-embedded tissue^5,6,16,17^. The ClearSee clearing protocol further improved optical penetration of root tissue and, when combined with fluorescent histochemical dyes, has revealed a rich spectrum of these structures, though so far in only a small number of species^16,18,19^. Together, these methods make it straightforward to generate large image collections spanning cultivars, species, genotypes, and conditions. Quantitative analysis of those images, however, has become the principal bottleneck limiting the scale and scope of biological study.

Quantification of root barriers has traditionally relied on categorical scoring^2,3^ or manual annotation^17^, approaches that are time-intensive and difficult to scale, especially for quantitative- genetic studies. The challenge is most acute for anatomically complex structures such as cortical aerenchyma, whose boundaries are often irregular, diffuse, or discontinuous, requiring complex methods, or making annotation subjective and prone to inter- and intra-annotator variability^15,20^. As a result, analyses that demand months of effort are typically confined to small sample sizes, and large image collections remain underused.

From a computer-vision standpoint, root anatomy poses a challenging structured-segmentation problem: the barriers and aerenchyma are not isolated objects but spatially organized, nested structures whose geometry and relative position are biologically constrained (**Fig. 1a**). Their morphology varies substantially across species, growth conditions, developmental stages, root types and further variability is introduced by staining intensity, sectioning quality, and imaging platform. Together, these factors make root anatomical segmentation a demanding, real-world task that goes well beyond controlled laboratory benchmarks.

**Figure 1:**
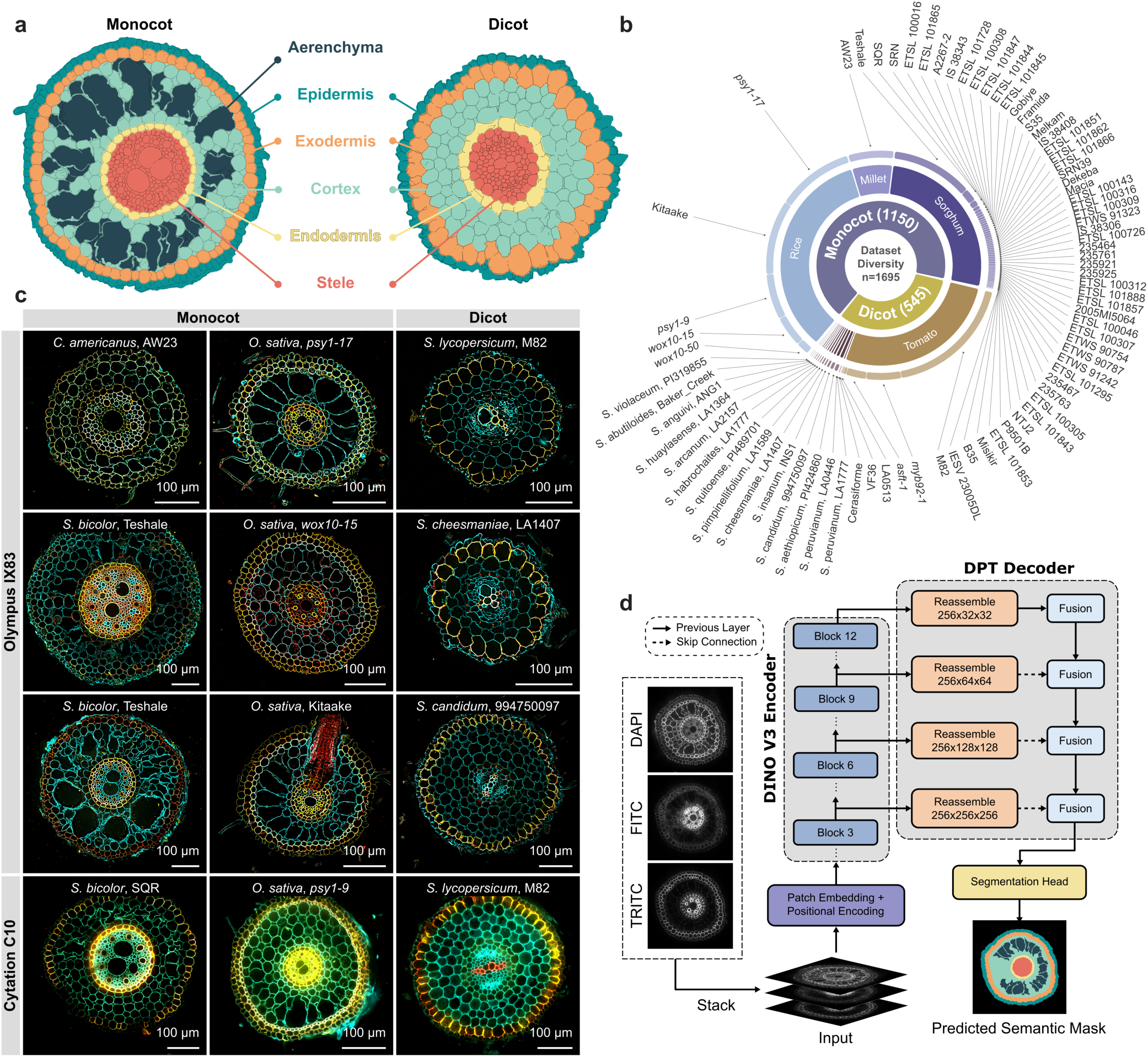
Anatomical structure segmentation pipeline overview. **(a)** Schematic of a root cross-section showing the six target anatomical classes: epidermis (teal), exodermis (orange), cortex (light teal), aerenchyma (dark teal), endodermis (yellow), and stele (salmon). Aerenchyma (air-filled cavities) are present only in monocot roots. **(b)** Species and genotype diversity of the full dataset. Sunburst chart of all 1,695 samples broken down by monocot versus dicot, then species (rice, millet, sorghum, and Solanum species), then genotype. Arc width is proportional to sample count, and each species is shown in the canonical color reused throughout the paper (rice, light blue; millet, light purple; sorghum, dark purple; Solanum species, tan). Train, validation, and test sets are detailed in **Supplementary Data 1 and Supplementary Table 1**. **(c)** Training data gallery. A 4x3 grid of training samples rendered as composite images (DAPI, cyan; FITC, yellow; TRITC, red). Rows 1-3 were imaged on the Olympus IX83 microscope (columns: millet/sorghum, rice, Solanum); row 4 was imaged on the Cytation C10 plate reader (sorghum, rice, Solanum). **(d)** RADIX (Root Anatomy Deep-learning Image Segmentation across Species and Platforms), our semantic segmentation model for mapping root anatomy. For each sample, the three fluorescence channels (DAPI, FITC, TRITC) are stacked as the model input (**Methods**). The image stack is passed through a vision-transformer encoder (DINOv3-S/16) and a dense-prediction-transformer (DPT) decoder, which together produce a per-pixel anatomical-class prediction. Left to right: the three raw channels, the composite, and the model’s prediction rendered with the same anatomical-class palette as in (a).

Machine-learning segmentation offers a route past these limitations, enabling rapid, consistent, and scalable analysis of complex anatomy. Once trained, a model applies uniform decision criteria across large datasets, compressing analysis from months to minutes and enabling experimental scales that are impractical by hand. Critically, such models can support reproducible quantification across biological and technical conditions, but only if they are trained and evaluated on sufficiently heterogeneous data. Large-scale analysis of root anatomically complex differentiation features across species has been limited by the absence of large, expert-annotated datasets and standardized benchmarks. Existing tools are also mismatched to the problem: cell segmentation methods such as Cellpose^21^ target individual cells rather than tissue-level topology (cell wall boundaries, aerenchyma cavities, and concentric cell-type cylinders with irregular borders) and platforms such as MorphoGraphX^22,23^ are best used with transgenic boundary markers in crops and are not designed to quantify barrier composition (lignin or suberin). Emerging spatially resolved -omic approaches are typically restricted to specific imaging modalities, species, or reconstruction frameworks and do not provide large-scale, tissue-level segmentation benchmarks across diverse conditions. Compared with the medical- and natural-image domains, plant anatomical images that capture this real-world biological and technical variability remain markedly underrepresented in public resources.

Here we address this gap with the first large-scale expert-annotated benchmark of root anatomical structures at scale, comprising 1,695 fluorescence images of root cross-sections drawn from globally important crops spanning the monocot-dicot divide and exhibiting substantial diversity in anatomical feature formation, focusing on the cell wall barriers of the endodermis, exodermis and on cortical aerenchyma. The images cover six anatomical classes across 17 species and three imaging platforms (**Fig. 1b**). Annotation quality was treated as a primary design goal. Every image was labeled by professional annotators and curated by domain experts. The benchmark is deliberately designed to vary along three axes (genotype, species and imaging hardware) that determine whether a model transfers beyond the laboratory in which it was trained.

Research has shown that pre-training on a large and diverse image data yields representations that transfer across many unrelated tasks from few labels, and that degrade far less on data unlike the training set than those of models fit to a single curated dataset^24^. Rather than train a model from scratch on root cross-section images alone, we begin from DINOv3^25^, a large vision model that has already learned general-purpose visual representations, such as shapes, edges, textures, and object boundaries, from billions of natural images without human labels. Using our high-quality benchmark, we develop **RADIX**, or Root Anatomy Deep-learning Image Segmentation across Species and Platforms, by fine-tuning DINOv3 pre-trained encoder together with a dense-prediction-transformer decoder. RADIX segments all six structures at an accuracy approaching inter-annotator agreement and generalizes to unseen species, genotypes, growth conditions, and images from another laboratory and microscope system, outperforming models pre-trained on smaller image collections when tested on out-of-distribution images. We also found that a single unified model matches or exceeds monocot- and dicot- specialist models without sacrificing in-group accuracy, and its predicted masks yield downstream trait measurements that agree closely with expert annotation. Applying this framework across genetic backgrounds, environmental perturbations, and species, we find that root barrier traits are multidimensional and highly context-dependent, shaped by the interaction of genetic and environmental factors. This work establishes a scalable approach to dissect how root barrier architecture contributes to plant performance under environmental and biotic stress.

## Results

### A high-quality multi-species plant root fluorescence microscopy dataset annotated with six anatomical structures

Supervised deep-learning models learn to segment images from labeled examples, so their accuracy is bounded by the scale and diversity of expert-annotated training data and, critically, by how closely that data resembles the images the model will encounter. Most existing fluorescence microscopy datasets are restricted to a single species, imaging platform, or laboratory. Models trained on them tend to generalize poorly, i.e., they lose accuracy when applied to images that differ from the training set, forcing each new study to annotate fresh data and retrain a model from scratch ^21,26–28^. To overcome this bottleneck, we assembled a dataset of 1,695 root cross-section images spanning seventeen species and three fluorescence imaging platforms, each annotated by plant biologists for six anatomical structure classes (**Fig. 1a**). The dataset was designed to vary along three axes: genotype, species and imaging hardware, which determine whether a trained model can be deployed beyond the laboratory in which it was built.

The dataset spans the major evolutionary split in flowering plants: three monocot species (rice, sorghum, and pearl millet) comprise 1,150 samples, and fourteen dicot species (all in the genus *Solanum*) comprise 545 samples. Monocot and dicot roots differ substantially in cell-wall thickness, root diameter, cortical cell number and the presence of aerenchyma, so training on both forces the model to learn features that hold across divergent root forms rather than ones tied to a single organ plan. Within each species, samples span multiple genotypes (distinct genetic backgrounds) contributed by laboratories at several institutions and geographic regions (**Fig. 1b, Supplementary Fig. 1, Supplementary Data 1 and Supplementary Table 1**). This diversity is essential: a model trained on a single genotype tends to overfit, latching onto that genotype’s idiosyncratic anatomy instead of the general structure, whereas a model useful to the wider root biology community should recognize the same anatomical structure across the varied root morphologies of many genotypes.

Every root cross-section was stained with three fluorescent dyes, each labeling a distinct cell- wall component. Calcofluor White provides a general cell wall (cellulose) contrast, Fluorol Yellow 088 labels suberin, and Basic Fuchsin labels lignin; suberin and lignin being the two hydrophobic polymers that build the root’s intercellular diffusion barriers. The three dyes were imaged through the DAPI, FITC, and TRITC filter channels, respectively, and are displayed throughout as a cyan / yellow / red composite (**Fig. 1c, Supplementary Fig. 1**). To capture the optical and noise differences introduced by different microscopes, samples were imaged on three platforms: the Olympus IX83 microscope (1,501 samples), the Cytation C10 plate-reader imager (159 samples), and the Zeiss LSM 880 confocal microscope (35 rice samples). In practice, research groups image their sections on whatever microscope is locally available, therefore bundling three platforms into one training set yields a single model robust to this variation, rather than requiring each user to retrain on their own hardware.

Each sample was annotated with hand-drawn polygons by trained annotators on the Labellerr online platform^29^ then verified by expert biologists in custom annotation software (**Supplementary Fig. 2**). Annotation averaged approximately one hour of manual work per image, representing over 1,700 hours of combined annotator and expert time across the benchmark. Polygons followed a cell wall-inclusive convention: each ring-shaped tissue (epidermis, exodermis, and endodermis) was outlined along the outer edge of its cell wall, with one outer and one inner contour per ring, so that the same boundary rule applied across annotators, samples, and platforms. Across 1,695 samples, annotators drew 31,594 polygons: 1695 outer epidermal ring contours, 3,390 endodermal (one outer and one inner per sample) and 3,390 exodermal ring contours, and 23,119 aerenchyma polygons. Aerenchyma counts ranged from zero in dicot sections (which lack the feature) to dozens of separate cavities in mature monocot sections. Together, these polygons define the six anatomical classes the model is trained to predict: epidermis, cortex, endodermis, exodermis, stele, and aerenchyma. This is, to our knowledge, the most extensive annotated fluorescence microscopy dataset of plant root cross-sections, and the only one labeling multiple anatomical class across monocot and dicot species at the level needed for quantitative trait analysis rather than simple cell boundary detection.

We partitioned the data into the standard machine-learning splits (data subsets). Samples from the Olympus IX83 and Cytation C10 platforms were divided roughly 80/10/10 into a training set of 1,293 samples (used to fit the model’s parameters; **Supplementary Fig. 1, Supplementary Data 1**), a validation set of 182 samples (used to tune settings and decide when to stop training; **Supplementary Data 1**), and an in-distribution test set of 185 samples (held out entirely for final evaluation). The 35 Zeiss LSM 880 images, contributed by an independent laboratory/institution were withheld as a separate out-of-distribution test set (meaning the model saw no images from this platform during training; **Supplementary Table 1**). To prevent information leakage (the model seeing near-identical images, such as adjacent sections of the same root, in more than one subset), all samples from a given biological experiment were kept within a single split.

These two test sets bracket the failure modes that matter for real-world deployment. The in-distribution test set contains cross-sections from the same species and platforms as the training data, but from entirely separate biological experiments. It measures whether the model has learned the underlying root anatomy rather than memorizing experiment-specific cues (e.g., a particular specimen or a sectioning artifact) and it estimates the accuracy a researcher should expect when applying the model to new samples under their existing imaging conditions. The out-of-distribution test set, by contrast, consists of rice samples imaged on a microscope the model never saw during training (the Zeiss LSM 880) and collected from crown roots, a root with a distinct lineage compared to other root types, from the training and validation data. It simulates the most demanding scenario (a different laboratory adopting the model on its own hardware) and quantifies robustness to shifts in imaging platform, sample preparation, root type, and operator practice that lie beyond the diversity sampled during training. Together, the two sets probe distinct generalization regimes: the in-distribution set reveals whether the model has overfit to specific experiments in our lab, and the out-of-distribution set reveals whether it has overfit to the specific imaging platforms it has seen.

### RADIX matches the inter-annotator agreement ceiling and generalizes across species, microscopes, and an independent laboratory

Building on this dataset, we developed RADIX, a semantic-segmentation model that assigns every pixel of an image to one of a fixed set of classes, here the six root anatomical structures plus background. RADIX takes a three-channel fluorescence image as input and returns a per-pixel anatomical-class map. The model comprises two main components (**Fig. 1d**). The *encoder* is a vision transformer initialized from DINOv3, a “foundation model” that Meta pretrained on ∼1.7 billion natural images without manual labels^25^. It compresses the input into a learned feature representation capturing both local structure and the global organization of the image data. The *decoder* is a dense-prediction transformer (DPT)^30^ that expands this representation back to full resolution by fusing encoder features from multiple depths, combining fine boundaries with large-scale context in the final map (full architecture in **Methods**). Starting from the pretrained encoder, we fine-tuned it and trained the decoder from scratch, jointly, on the annotated training set (**Supplementary Fig. 1, Supplementary Data 1**), using percentile normalization and random augmentation to build robustness to imaging differences while preserving overall root morphology (**Methods, Supplementary Fig. 3**).

We applied RADIX to both test sets with an identical preprocessing pipeline and obtained consistently high accuracy across all anatomical structures. Predicted maps were nearly indistinguishable from expert annotations, resolving the relatively thin endodermal and exodermal rings and the irregular aerenchyma lacunae across all monocot species (**Fig. 2a**). We quantified accuracy with the intersection-over-union (IoU), the area where the predicted and ground-truth regions overlap divided by the area they jointly cover, ranging from 0 (no overlap) to 1 (perfect overlap), and report the mean across the six classes as mIoU (**Methods**). On the in-distribution test set, the stele and cortex each exceeded IoU 0.970, while the endodermis, exodermis, epidermis, and aerenchyma reached 0.912, 0.878, 0.820, and 0.601, respectively (**Fig. 2a,b; Supplementary Table 2**). Accuracy held on the out-of-distribution (Zeiss) test set: stele, cortex, exodermis, endodermis, and aerenchyma reached 0.974, 0.982, 0.847, 0.858, and 0.835, and epidermis, 0.738 (**Fig. 2a,c; Supplementary Table 2**). RADIX also handled the disruption of radial symmetry resulting from emerging lateral roots: the *Solanum lycopersicum* cv. M82 and *Oryza sativa* Kitaake samples in **Fig. 2a** contain lateral roots that break the normal radial symmetry, as indicated by the arrowhead, yet all structures were segmented correctly (mIoU 0.913 and 0.869).

**Figure 2:**
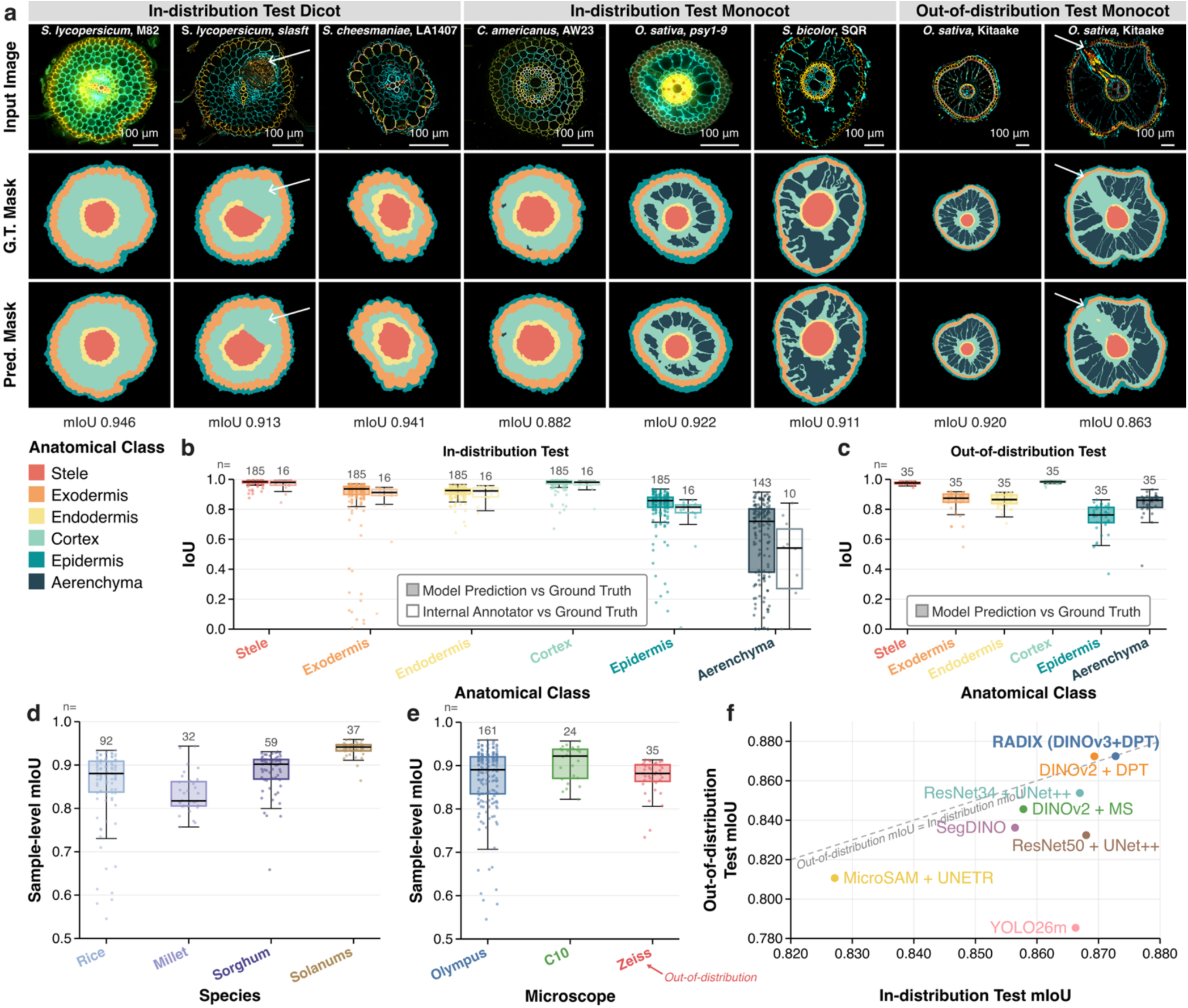
RADIX segmentation accuracy approaches the inter-annotator agreement ceiling and generalizes across species, microscopes, and an independent lab. **(a)** Example RADIX predictions on representative samples from the in-distribution and out-of-distribution test sets. For each sample, the input stack (DAPI, cyan; FITC, yellow; TRITC, red), ground-truth mask, and predicted mask are shown side by side. Sample-level mIoU across the six anatomical classes is reported below each prediction. Arrows indicate lateral root primordia. **(b)** Per-class IoU on the in-distribution test set. Filled boxes show IoU between RADIX predictions and ground-truth annotations; box plots display the median, interquartile range (IQR), and 1.5xIQR whiskers, with individual samples overlaid as jittered points. Open boxes show IoU between a second internal annotator and the ground-truth annotation on 16 samples, establishing a per-class inter-annotator agreement ceiling. RADIX’s per-class IoU sits at or near this ceiling for every class. **(c)** Per-class IoU on the out-of-distribution test set. Box plot conventions as in (b). **(d)** Sample-level mIoU stratified by species. Rice pools both the in- and out-of-distribution test sets; other species are from the in-distribution test set only. **(e)** Sample-level mIoU stratified by microscope (Olympus, C10, Zeiss). Zeiss is the out-of-distribution platform. **(f)** In-distribution vs. out-of-distribution mIoU for RADIX and seven alternative segmentation architectures, all trained and evaluated on identical data sets. Each point represents one architecture; the grey dashed line marks y = x, where out-of-distribution performance equals in-distribution performance.

The lower IoU values for aerenchyma and epidermis stem from the intrinsic geometry of these structures or their difficulty in sample preparation, respectively, not from segmentation failure. Aerenchyma consists of sparse, micron-scale cavities bounded by cell wall remnants only one to two pixels wide. The epidermis is a thin outer layer, which can be compressed or damaged during sample embedding and sectioning, and often stained weakly (**Fig. 2a, Supplementary Fig. 4**). Because IoU is a ratio of overlapping area to total area, a small, fixed boundary misalignment removes a disproportionately large fraction of a thin object’s area, so the metric understates the true visual agreement for these classes (**Supplementary Fig. 4**). The same effect explains the apparent species-level differences. RADIX achieved mean IoU of 0.883, 0.843, 0.834, and 0.936 on sorghum, rice, pearl millet, and Solanum, respectively (**Fig. 2d, Supplementary Table 3**); pearl millet’s slightly lower score is a direct consequence of its aerenchyma being roughly an order of magnitude smaller than in rice or sorghum, which makes per-pixel IoU especially sensitive to minor boundary shifts despite visually accurate masks (**Fig. 2a**). Consistent with this, aerenchyma IoU is *higher* on the out-of-distribution rice set (0.835) than in-distribution (0.601), because those sections contain larger lacunae, further evidence that the score tracks object size rather than segmentation quality.

To test whether the residual error reflects intrinsic annotation ambiguity rather than a model limitation, we performed an inter-annotator comparison, which sets a practical ceiling: a model cannot be expected to agree with the ground truth more closely than two human experts agree with each other. A second expert plant biologist re-annotated a balanced subset of 16 samples spanning monocot and dicot species, blinded to the consensus ground truth. This annotator agreed with the consensus at mean IoU 0.856, whereas RADIX reached 0.893, matching or exceeding the human on every anatomical class (**Fig. 2b, Supplementary Table 4, Supplementary Fig. 5**). The largest human-model gaps were on aerenchyma (model 0.518, human 0.455), epidermis (0.830 vs. 0.747), and endodermis (0.922 vs. 0.870), classes whose boundaries hinge on subjective interpretation of weakly stained or developmentally variable features. Notably, aerenchyma was the lowest-scoring class for both the model and the human, indicating that its residual error reflects irreducible biological and imaging ambiguity rather than a model deficiency. RADIX has therefore reached the practical upper bound achievable given the inherent ambiguity in these images.

Finally, RADIX performed consistently across imaging platforms. On the in-distribution test set it achieved per-platform mean IoU of 0.905 on the Cytation C10 plate-reader imager and 0.868 on the Olympus IX83 microscope (**Fig. 2e).** On the out-of-distribution test set, 35 rice samples at an unseen mature root of a different type, imaged on a Zeiss microscope absent from training, it achieved a nearly identical mIoU of 0.878, showing that the model transfers robustly to data collected by an independent laboratory on unfamiliar hardware.

### RADIX outperforms other root anatomical segmentation models

To understand what makes the RADIX segmentation model robust, we trained seven alternative ML architectures on the same data under an identical loss, augmentation pipeline, optimizer, and schedule (**Methods, Supplementary Fig. 6a**). We chose them to span the design space along three axes: the encoder’s pre-training, the decoder, and the prediction paradigm (**Supplementary Table 5**). Two are convolutional baselines: ResNet34 and ResNet50 encoders pretrained on ImageNet^31,32^, paired with a UNet++ decoder^33^, the standard convolutional, skip-connection approach used in most prior plant-image segmentation^33^. Three probe RADIX’s own design: one keeps the DPT decoder but swaps the DINOv3 encoder for the earlier, smaller-data DINOv2^34^ (isolating the encoder generation), and two keep a DINO encoder but replace the DPT decoder with simpler prediction heads, a multi-scale linear head with DINOv2, and SegDINO^35^, a lightweight per-patch multilayer perceptron with DINOv3 (isolating the decoder). The sixth pairs MicroSAM^36^, a microscopy-tuned version of Meta’s Segment Anything Model (SAM)^37^, a foundation model pretrained directly on segmentation rather than on generic images, with a transformer UNETR decoder^38^. The seventh, YOLO26m-seg^39^, is a state-of-the-art *instance*- segmentation model (it detects and outlines individual objects rather than labeling every pixel) pretrained on COCO^40^, representing the per-object detection paradigm common in off-the-shelf segmentation packages^41,42^. Among the models evaluated, RADIX is fine-tuned from the encoder with the broadest pre-training.

In-distribution, all eight models scored within a narrow band of mIoU 0.827-0.873 (**Fig. 2f, Supplementary Table 5**), showing that the dataset is rich enough for several modern ML architectures to fit the training distribution, and that in-distribution accuracy alone does not distinguish them. The architectures separated only under domain shift. On the out-of-distribution Zeiss set the spread widened to 0.785-0.872, and that spread tracks two design choices. First, the encoder’s pretraining: the self-supervised DINO encoders held up best (RADIX/DINOv3+DPT and DINOv2+DPT, both 0.872), outperforming the ImageNet-pretrained convolutional ResNet (0.854 ResNet34; 0.832 ResNet50) and, notably, the segmentation- pretrained SAM encoder (MicroSAM+UNETR, 0.811), a generic self-supervised prior transferred across imaging platforms better than one pretrained directly on segmentation. Second, the decoder: holding the encoder fixed, the multi-scale DPT decoder (0.872) beat the simpler heads (0.846 DINOv2 + multi-scale-linear; 0.836 SegDINO), indicating that multi-scale feature fusion helps preserve thin, nested boundaries under shift. The instance-detection model degraded most (YOLO26m-seg, 0.785), consistent with nested ring-within-ring tissue being a poor fit for box-based object detection. Most models fall below the grey dashed y = x line in **Fig. 2f**, they lose accuracy when transferred to an unseen platform, whereas RADIX sits at the top on both axes. RADIX (DINOv3 + DPT) thus maximizes both robustness-determining factors and achieves the best in-distribution and out-of-distribution performance of all architectures tested, while using fewer parameters than the ResNet50 and SAM baselines, establishing it as a robust default for root-anatomy segmentation.

### Unified RADIX model for monocots and dicots generalizes better than the specialist models trained on monocots or dicots individually

Monocot and dicot root cross-sections differ substantially, in cell-wall thickness, root diameter, the number of cell layers, and the presence of aerenchyma. A natural concern with training one model jointly on both is negative transfer, i.e., the model might dilute its representation of one group to accommodate the other, so that a “specialist” trained on a single species would beat the joint “generalist” on that group’s own samples. To test this directly, we trained two additional RADIX models with an identical architecture, augmentation, and training pipeline to the unified model, differing only in training-set composition. The Monocot Specialist saw only the 856 monocot training samples (rice, sorghum, and pearl millet); the Dicot Specialist saw only the 437 *Solanum* training samples; and the unified model used in above was trained on all 1,293 samples together. We split the in-distribution test set by lineage (monocot vs. dicot) and evaluated all three models on 148 monocot in-distribution samples, 37 dicot in-distribution samples, and the 35 out-of-distribution Zeiss rice samples (**Fig. 3a**).

**Figure 3:**
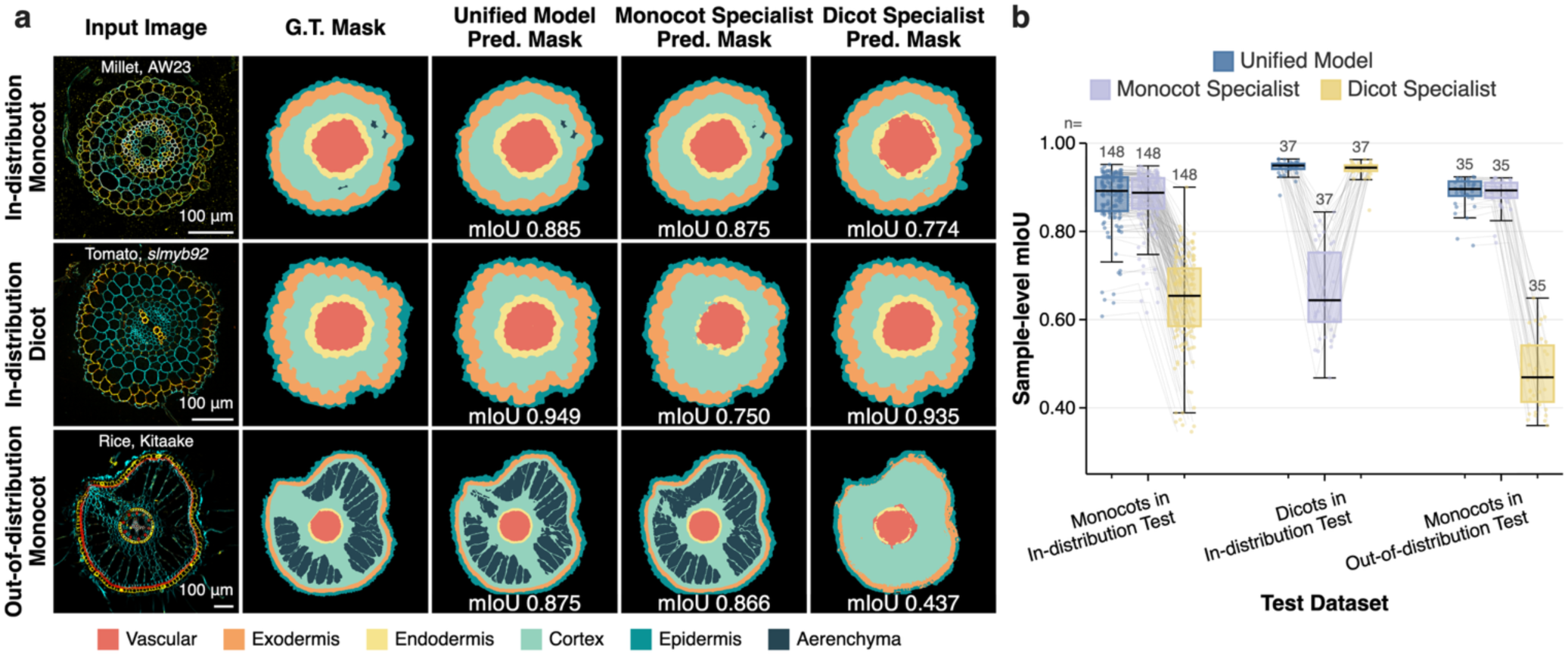
A unified RADIX model trained on both monocots and dicots generalizes better than specialist models. **(a)** Example segmentation predictions on three representative samples (one in-distribution monocot, one in-distribution dicot, and one out-of-distribution monocot for three models. The Unified Model was trained on all species, the Monocot Specialist on rice, millet, and sorghum only, and the Dicot Specialist on Solanum species only. For each sample, the input stack (DAPI, cyan; FITC, yellow; TRITC, red) is shown alongside the ground-truth annotation and the three model predictions. Specialists fail on out-of-distribution inputs. The Monocot Specialist underperforms in segmenting endodermis and exodermis on the tomato cross-section, and the Dicot Specialist mislabels aerenchyma, endodermis, and exodermis on both monocot samples. The Unified Model recovers all structures with high IoU in all three cases. **(b)** Per-sample mIoU (across the six anatomical classes) for the three models on three test sets (in-distribution monocot test, in-distribution dicot test, and monocot out-of-distribution). Box plots show the median, IQR, and 1.5xIQR whiskers, with individual samples overlaid as jittered points. Thin grey lines link each sample’s mIoU across the three models within a set, allowing within-sample shifts between Unified and Specialist predictions to be read directly. The Unified Model matches or exceeds the in-group Specialist on each group’s own test set and substantially outperforms the off-group Specialist on every test set, demonstrating that mixing monocot and dicot samples during training does not sacrifice in-group accuracy while recovering most of the cross-group generalization.

Within its own training group, each specialist was matched by the unified model. On the 148 monocot in-distribution samples, the Monocot Specialist and the unified model reached mIoU (averaged over the six anatomical classes) of 0.856 and 0.857; on the 37 dicot in-distribution samples, the Dicot Specialist and unified model reached 0.932 and 0.936; and on the 35 Zeiss out-of-distribution rice samples, the Monocot Specialist and unified model reached 0.872 and 0.873 (**Fig. 3b**). Joint training across both clades therefore preserves full specialist-level accuracy on each group rather than diluting it, at no measurable cost.

By contrast, performance collapsed when each specialist was applied to the group it had never seen during training. The Monocot Specialist fell from mIoU 0.856 on its own test set to 0.603 on dicots, a drop of 0.253. The Dicot Specialist fell from 0.932 to 0.589 on monocots (a drop of 0.343), and further to 0.401 on the out-of-distribution samples (a drop of 0.531 from its dicot accuracy), the last compounding two shifts at once, an unseen group and an unseen imaging platform. The unified model, by contrast, stayed between mIoU 0.857 and 0.936 across every test set. Representative predictions (**Fig. 3a**) show the failure directly: the off-group specialists either miss entire structures or mislabel large regions as the wrong class, whereas the unified model’s predictions remain essentially indistinguishable from the expert ground truth on both monocot and dicot samples.

### RADIX’s high segmentation performance translates to high accuracy in various downstream quantification tasks

Segmentation masks become scientifically useful only when they translate into accurate biological readouts, whether quantitative measurements, spatial analyses, or cell-type-resolved molecular profiling. Having established that RADIX delineates root anatomical structures at or even better than human-annotator quality, we next demonstrate how predicted masks can be directly used for various downstream trait quantification.

Within each predicted mask, fluorescence intensities are thresholded to exclude low-intensity pixel contributions from within the cell, isolating the signal from the cell wall itself. FITC and TRITC channel intensities serve as proxies for suberin and lignin deposition, respectively, as the fluorescent dyes used for histochemical staining (Fluorol Yellow 088 and Basic Fuchsin) are imaged in these channels ^16^. Mean FITC and TRITC intensities within the predicted endodermis and exodermis masks therefore reflect cell-wall biochemical composition rather than being confounded by cell size (**Fig. 4a-d**).

**Figure 4:**
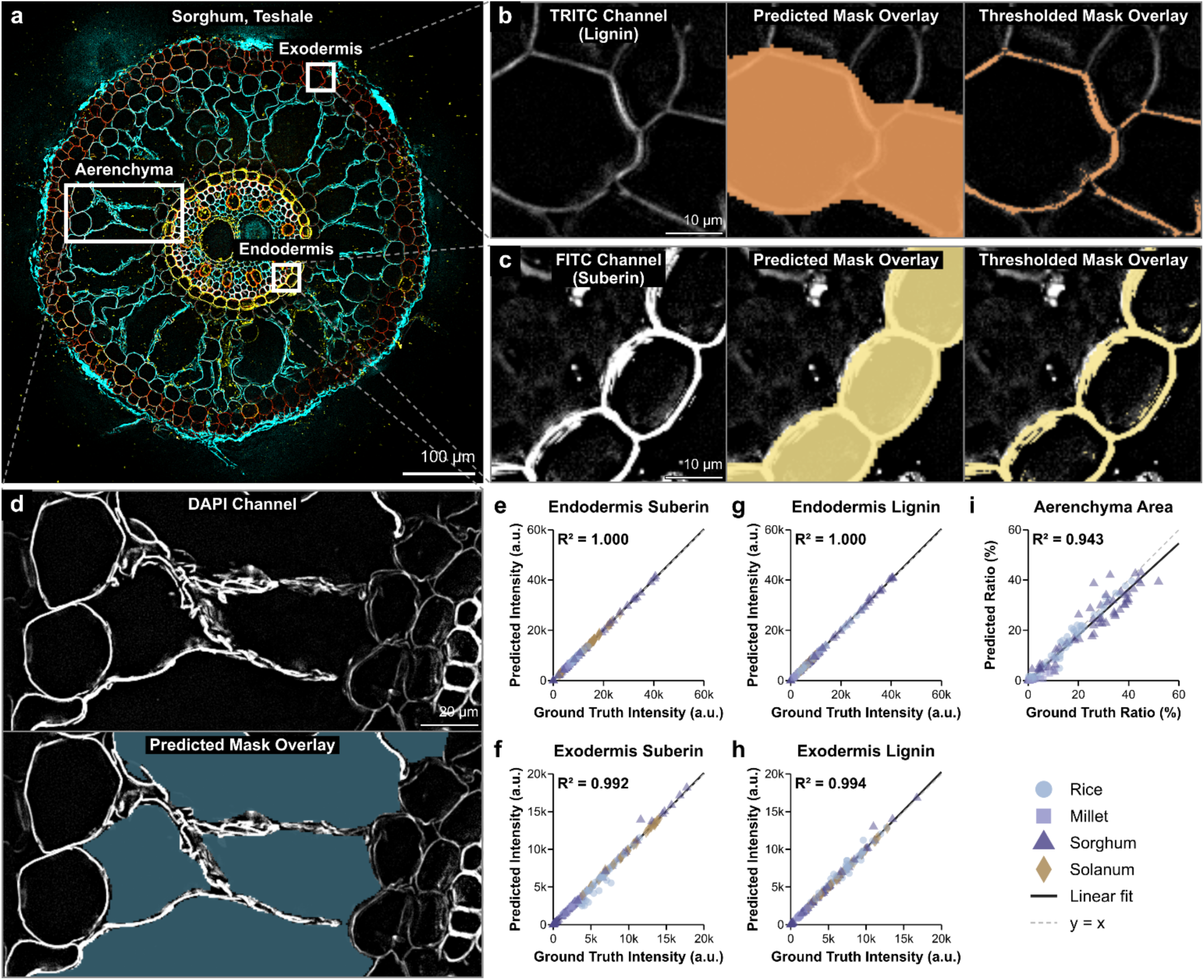
RADIX’s segmentation accuracy translates to accurate downstream quantification of root anatomical traits. **(a)** Example image stack of DAPI (cyan), FITC (yellow), and TRITC (red) channels for a Sorghum N3 genotype imaged on an Olympus microscope. Dashed lines link the endodermis, exodermis, and aerenchyma crop boxes to their zoomed views in (b), (c), and (d). **(b)** Exodermis zoom-in showing, left to right, the FITC channel (suberin), the predicted mask overlay, and the overlay restricted to pixels above a suberin intensity threshold. **(c)** Endodermis zoom-in showing, left to right, the TRITC channel (lignin), the predicted mask overlay, and the overlay restricted to pixels above a Casparian-strip lignin threshold. **(d)** Aerenchyma zoom-in showing the DAPI channel (top) and the predicted aerenchyma mask overlaid on DAPI (bottom). The prediction cleanly follows the inner cell-wall border of the air-filled cavities. **(e-h)** Predicted vs. ground-truth mean fluorescence intensity within the thresholded predicted mask, for endodermis suberin (e, FITC), exodermis suberin (f, FITC), endodermis lignin (g, TRITC), and exodermis lignin (h, TRITC), across all held-out test samples (n = 185 from Rice, Millet, Sorghum, and Solanum). **(i)** Predicted vs. ground-truth aerenchyma area ratio (aerenchyma area divided by total root area, expressed as a percentage) on the same test set, restricted to monocot samples (n = 148). Solanum samples are excluded because they lack aerenchyma. In each scatter plot (e to i), the solid black line is the per-panel linear regression and the dashed grey line marks y = x. The top-left annotation reports R², the fitted equation, and the sample count. Marker shape and color encode species (rice, millet, sorghum, Solanum), using the same palette as Fig. 1b.

To validate this approach, we compared trait values extracted from predicted masks against those extracted from expert-annotated ground-truth masks across all 185 held-out test samples spanning rice, millet, sorghum, and tomato. We quantified FITC and TRITC intensities within the predicted endodermis and exodermis masks, reflecting suberin and lignin deposition in each cell wall and cell type, respectively. Predicted and ground-truth fluorescence intensities agreed with near-perfect correlation across all four traits (R² = 0.992-1.000; **Fig. 4e-h**; n = 185 monocot and dicot samples), with regression slopes close to unity, indicating no systematic bias introduced by mask error. Aerenchyma area ratio, the fraction of total root cross-sectional area occupied by aerenchyma, showed equally high agreement between predicted and ground-truth measurements (R² = 0.943; **Fig. 4i**; n = 148 monocot samples).

The full pipeline, including segmentation and batch thresholded intensity extraction, processes a complete experimental set at ∼1.2 s per image on a consumer GPU (NVIDIA GeForce RTX 3070 Ti) and ∼3.8 s per image on a CPU-only workstation (AMD Ryzen 9 7900X), with fluorescence thresholds defined once per experimental batch rather than adjusted per image. At this throughput, 100 images are processed in 2 min on GPU and under 7 min without a dedicated GPU. This represents a 1,500- to 3,000-fold improvement over the 30-60 min required for expert manual annotation of a single cross-section depending on anatomical complexity on GPU, and a 500- to 950-fold improvement on CPU, making large-scale quantitative studies tractable without specialized computing infrastructure, reducing what would otherwise require weeks of expert time for a typical experiment to a fully automated analysis completed in minutes.

Together, these results confirm that RADIX’s segmentation accuracy translates directly into quantitatively reliable trait measurements, without the need for manual correction or post-hoc calibration, enabling its application to biologically diverse experimental contexts.

### Root cell type anatomical features are multi-dimensional and context dependent

We applied our automated segmentation framework to elucidate biologically meaningful insights across diverse contexts in both monocot and dicot species. These included a set of naturally varying local land races, released varieties and cultivars, a CRISPR-edited mutant line, and the influence of biotic and abiotic perturbation.

#### Natural genetic variation across sorghum genotypes

Ethiopian sorghum germplasm genetically varies in drought tolerance and resistance to the parasitic plant, *Striga hermonthica*^43,44^. Lignin and suberin are associated with these traits^5,17,45,46^ and to determine if biopolymer levels are candidates to underlie these traits, we quantified exodermal lignin and suberin in ten Ethiopian sorghum genotypes (**Supplementary Table 6**) and evaluated whether the model generalizes across their genetically variable backgrounds. Root cross-sections from ten-day old root plants were triple stained with calcofluor white, basic fuchsin and Fluorol Yellow. The model accurately segmented key anatomical features; including aerenchyma space percentages, endodermal and exodermal layers, and stelar tissues and revealed variation in lignin and suberin in both the endodermis and exodermis.

To the best of our knowledge, exodermal lignin and suberin have not been quantified in sorghum. Genotypes displaying high exodermal lignin and suberin are distinct from those displaying high endodermal lignin and suberin, suggesting distinct developmental regulation. A single genotype may have both high lignin and suberin, expressed to the relative lowest accumulating genotypes (**Fig. 5a**). Genotype 1 (Wetetbegunchie) has a 1.3- and 1.2-fold increase in endodermal lignin compared to genotypes 51 (ETSL 100576) and 56 (10 SR-RIBKA = 235921; p values < 0.05), and a 1.3-fold increase in endodermal suberin compared to genotype 51 (p values < 0.05) (**Fig. 5a**). Genotype 17 (Framida) has a relatively high abundance of exodermal lignin and suberin signals - 40% more for both (p values < 0.05) compared to genotype 56 (**Fig. 5a**). In contrast, some genotypes have either high endodermal lignin (genotype 50/ETSL 100726) or suberin (genotype 45/IS 38343). This was also observed for the exodermis where genotype 51 (ETSL100576) has only high exodermal suberin relative to other genotypes. These results demonstrate robust performance of the model across genetic backgrounds, its use in mapping loci associated with quantitative variation in these traits and their role in environmental adaptation. We next asked whether the model can resolve environmentally induced plasticity in these traits.

**Figure 5:**
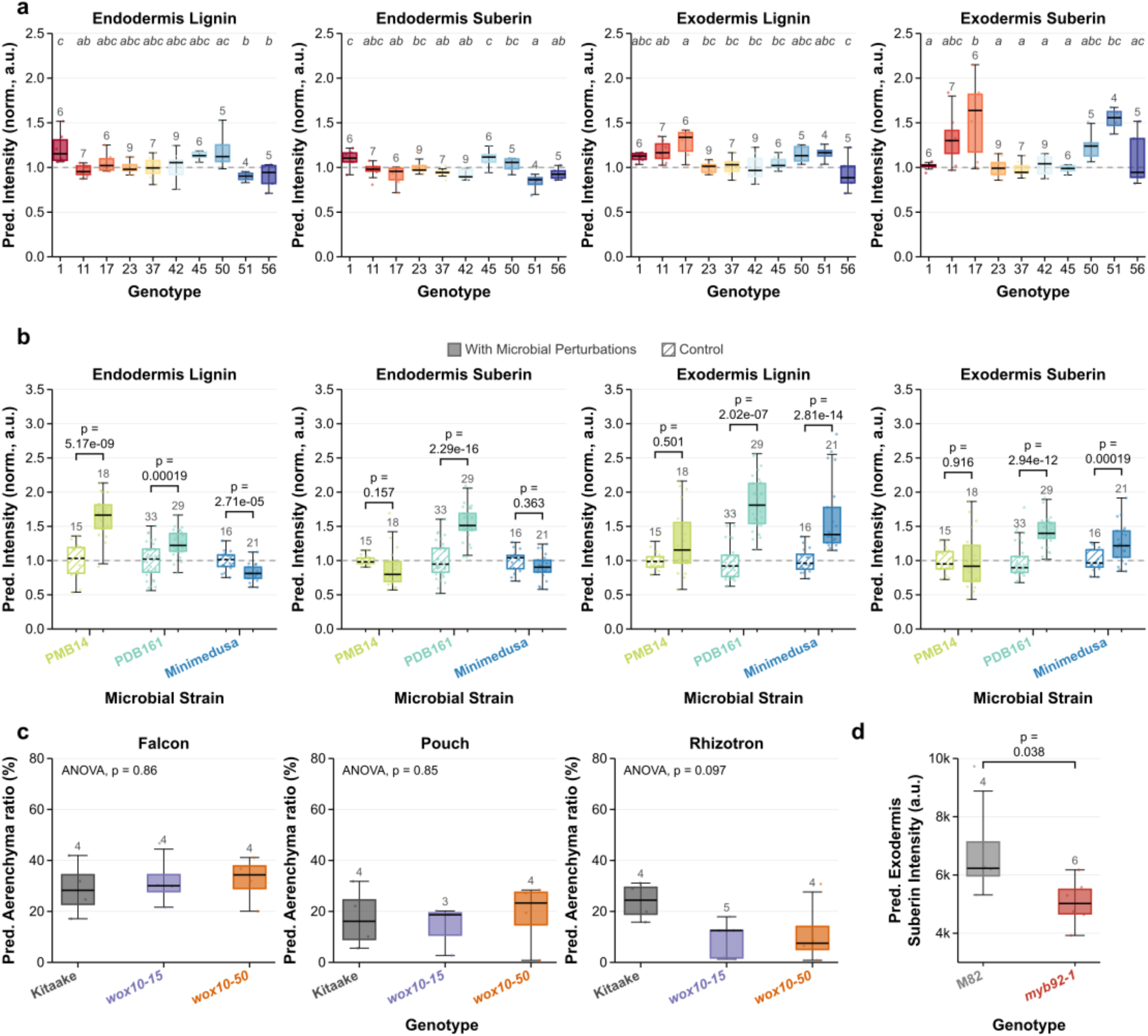
Automated segmentation quantifies root anatomical barrier variation across species, genotypes, microbial treatments, and growth systems. **(a)** Endodermal and exodermal barrier intensity across selected Sorghum bicolor genotypes. Fluorescence intensities of lignin (TRITC) and suberin (FITC) channels in the endodermis and exodermis are shown for 10 selected sorghum genotypes normalized to genotype 23 (Teshale). Compact letter displays (CLD) above each group indicate statistically distinct genotypes (one-way ANOVA with Tukey HSD post-hoc test, α = 0.05). Values are averaged per plant prior to statistical testing. **(b)** Microbial perturbations of root barrier traits in S. bicolor cv. Teshale. Barrier intensities are shown for three microbial inoculants: PDB161 (Pseudomonas uvaldeensis), Minimedusa (Minimedusa polyspora), PMB14 (Microbacterium foliorum), alongside within-experiment controls, drawn from a large multi-batch screening series. Exact adjusted p-values are shown above each comparison (linear mixed-effects model with Experiment as random effect; Benjamini-Hochberg FDR correction across all traits and treatments). Individual data points represent cross-sections; n = 5 biological replicates (plants) per treatment with 3-7 cross-sections per plant. **(c)** Aerenchyma ratio in Oryza sativa wox10 loss-of-function mutants across three growth systems. O. sativa (cv. Kitaake) plants and wox10-15 and wox10-50 loss-of-function mutant alleles (Kitaake background) were grown in Falcon tubes, paper pouches, and rhizotrons. Data are averaged per plant prior to statistical testing. Exact p-values are shown relative to wild type (one-way ANOVA). **(d)** Exodermal suberin (FITC) intensity in Solanum lycopersicum myb92-1 mutants. Exact p-value is shown relative to wild type (two-sided Mann-Whitney U test; n = 4 wild type, n = 6 myb92-1 plants). In all panels, boxes show median, interquartile range (IQR), whiskers extend to 1.5x IQR.

#### Microbial perturbations reshape barrier traits

Microbiome and microbial induction of endodermal suberin and lignin have previously been reported^11,17^ and root apoplastic barriers can coordinate spatial localization of bacteria^47^. To determine whether the model can resolve such environmental-induced plasticity in root cell type barriers, we analyzed sorghum roots exposed to three microbial isolates: *Pseudomonas uvaldensis* (PDB161), *Microbacterium foliorum* (PMB14), and *Minimedusa polyspora* (Minimedusa) that were entirely excluded from model training. Microbial inoculation of the Teshale sorghum genotype, subsequent sectioning, histochemical staining and application of RADIX, revealed diverse induction modes of cell type barriers. While one induced a uniform shift across all cell type barriers (PDB161), another induced a change in only a single cell type barrier (PMB14, endodermal lignin), while another reduced endodermal lignin and increased exodermal lignin (Minimedusa). Despite this biologically induced variation, the model consistently captured anatomical feature differences across microbial inoculants (**Fig. 5b**; **Supplementary Figure 7**), enabling quantitative comparison of inoculation-dependent responses^17^ and provides a framework to elucidate microbe-specific changes in cell type development that facilitates tolerance to environmental perturbations. Environmental interactions with root development are further dependent on genotype. We therefore next examined whether environmental plasticity can be mechanistically dissected using targeted genetic perturbation.

#### *WOX10* by environment-dependent control of rice aerenchyma formation

Aerenchyma formation is highly environmentally plastic and associated with drought and flooding tolerance^48–50^. We employed RADIX to characterize the root cellular phenotype of a mutant for a cell type-enriched gene, *WOX10* (*Os08g0242400*). Two independent CRISPR-Cas9-edited rice *wox10* mutant alleles (*wox10-15* and *wox10-50*), each carrying a unique 1-bp insertion in the open reading frame that is predicted to cause a frameshift mutation (**Supplementary Figure 8**). Root anatomical traits were compared between the wild-type and *wox10* mutants grown across multiple growth systems, including vermiculite-based rhizotrons that were entirely excluded from model training. An ANOVA indicates that the growth system is the most informative factor in aerenchyma formation (p-value = 0.0010). When grown in sand or paper pouches, the *wox10* mutant alleles exhibited aerenchyma formation comparable to the wild type (**Fig. 5c**). In contrast, when grown in rhizotrons, the mutant alleles had reduced aerenchyma relative to wild type (ANOVA, p-value <0.1). These differences were consistently captured through quantitative comparison by the model and manual annotation (Pearson correlation value r = 0.99, p-value < 0.0001). Together, these results support the environmental plasticity of aerenchyma formation and suggest that WOX10 plays a role in this variable feature.

Having demonstrated that condition-dependent effects on aerenchyma formation can be resolved in rice, we next asked whether the model can similarly capture a cell type-specific barrier phenotype in a dicot species.

#### Genetic regulation of exodermal suberization in tomato

To assess whether the model can resolve a cell type-specific cell wall barrier phenotype in a dicot species, we utilized a previously characterized *slmyb92-1* tomato transcription factor knockout mutant in which exodermal suberin was previously observed to be significantly reduced^5^ in a fully held-out experiment not included in model training. As with rice, thin cross-sections generated from primary roots of wild type tomato (cv. M82) and *slmyb92-1* were stained for suberin with Fluorol Yellow 088. Quantitative analysis revealed that the mutant exhibited reduced Fluorol Yellow fluorescence compared to wild type, in line with the published phenotype (**Fig. 5d**).

Together, these results demonstrate that root anatomical barrier traits are shaped by both genetic variation and environmental inputs, and that their interaction can be quantitatively resolved across species. By enabling consistent, cell-type-specific phenotyping across diverse experimental contexts, automated segmentation via RADIX provides a scalable framework for dissecting the genetic and environmental regulation of root adaptive cell type features.

## Discussion

### Model performance and generalization enable biological discovery

Rather than reporting a single accuracy number, we found that models that are indistinguishable on in-distribution data diverge sharply for out-of-distribution data. With RADIX leading on both data distributions, we attribute the good generalizability largely to the large-scale high-quality expert-annotated dataset for fine tuning and two design choices. The first is an encoder pre-trained on billions of image data, whose features transfer across imaging platforms better than alternatives pre- trained at smaller scale. The second is a multi-scale fusion decoder, which has shown to preserve the thin, nested boundaries that define root anatomy. Root cross-section images differ from one another in staining intensity, contrast, focus, sectioning artifacts, and scale, which is a subset of the variation spanned by billions of natural images. An encoder trained on them therefore arrives already partially invariant to the differences between microscopes and growth systems, without ever having seen a root. Fine-tuning can then spend a limited label budget on anatomy rather than on rediscovering edges and textures. This could also explain why the unified model outperforms the clade specialists. Images of any species inform the shared representation of a suberized wall or an aerenchyma cavity, and splitting the training set by clade discards that transfer while halving the data available to each model. The alternative models with smaller pre-training could underperform for the same reason. Encoders pre-trained on narrower datasets inherit a correspondingly narrower range of invariances, which suffices on in-distribution data and fails when imaging conditions move. This implies that performance under domain shift is set less by the network than by the breadth of what it saw before the task and the heterogeneity of the data used to adapt it.

Root anatomy is defined by thin, nested cell-wall boundaries that a coarse prediction would blur together. We found that using a decoder that fuses features across spatial scales lets the model recover detailed boundaries while still using the encoder’s high-level context to distinguish one tissue ring from another. Together these two design choices allow RADIX segmentation to be robust to laboratory-specific differences in sample preparation, microscope choice and availability. In plant biology, differences in root anatomical architecture between monocots and dicots reflect their evolutionary divide (140-200 million years ago), including variable cell layers and differentiation features, leading to tools or frameworks that could be applied to only a small number of species within each lineage, or even a genus. RADIX can be applied across all of these without per-dataset retraining, so that differences measured downstream reflect biology rather than the imaging platform or a model re-fit to each experiment.

This is a capability prior approaches did not provide: manual and categorical scoring at cell type- resolution do not easily scale to genome-wide association screens^11,51^; laser ablation tomography is accurate but instrument-limited^9,15^; and general cell-segmentation tools (e.g., Cellpose and MorphoGraphX) target individual cells or require boundary labels^21,22^, which in plants requires staining of the plant cell wall, or generation of transgenic lines with membrane markers. This limits the resolution of cell-type-level tissue topology and prevents its use in species that cannot be transformed, quantification of cell-wall biopolymer types and levels, and the “dead” space of aerenchyma. By validating against an inter-annotator ceiling and across a held-out imaging platform, we establish that the segmentation is reliable enough to serve as the measurement layer for quantitative root anatomy.

Root anatomical barriers or aerenchyma are typically manually analyzed one trait at a time, which makes it difficult to see how they vary together. By quantifying suberization, lignification, and aerenchyma simultaneously across genotypes and conditions, we observed that these features do not vary independently: they shift in trait-specific, context-dependent ways that are most naturally described multi-dimensional variation in barrier or anatomical architecture.

Capturing this structure requires measuring several anatomical features at once at scale, precisely what manual workflows make impractical, and the patterns we report here should be read as a demonstration of that capability across representative genetic and environmental contexts rather than an exhaustive survey.

A defining feature of this system is its pronounced environmental plasticity. Microbial treatments and growth conditions induced coordinated, or cell type/genotype-specific changes in barrier composition and structure, rather than uniform shifts across all features. Alterations in suberin deposition, lignification, and aerenchyma formation varied in both magnitude and direction depending on the microbial context, genotype or environmental stimulus, supporting prior observations, but increasing the spectrum of cell type changes possible to be modulated. These observations support a model in which root barrier architecture is dynamically adjusted in response to external cues, enabling plants to fine-tune transport properties and interactions with their surroundings. Beyond screening genotypes and environments at scale, accurate tissue- level segmentation is a prerequisite for placing molecular measurements in anatomical context. Spatial transcriptomics and single cell and single nucleus RNA sequencing increasingly resolve cell type-specific gene expression, but interpreting those data requires assigning signal to defined anatomical patterning or differentiation features, like cell-type cylinders, barriers, and aerenchyma that RADIX delineates. Coupling automated anatomical segmentation with spatial molecular profiling could link barrier composition to the regulatory programs that produce it, across the species and conditions in which these traits matter agronomically.

### Limitations and future directions

Despite these advances, several considerations remain. The framework operates on static cross-sectional images and therefore depends on experimental design to capture specific biological processes. While our dataset includes sections from different root types, resolving temporal dynamics of barrier formation requires dedicated time- course experiments or longitudinal sampling strategies. In addition, segmentation enables precise quantification of structural and compositional features, but interpretation of these traits in a functional (adaptive) context depends on complementary experimental approaches to validate including generation of mutant backgrounds lacking or producing excess of these cell type features. The framework is therefore best viewed as a quantitative phenotyping tool that can be integrated with assays probing physiological consequences of barrier variation. Model performance may also be influenced by annotation quality and dataset composition, particularly in cases of ambiguous boundaries or low signal-to-noise ratios. Expanding training datasets to include additional species, imaging conditions, and challenging anatomical cases will further improve robustness and generalization. More broadly, the ability to extract consistent, cell-type- specific anatomical measurements at scale creates new opportunities for integrating image- based phenotyping with experimental designs targeting developmental progression, genetic perturbation, and environmental responses. This approach may also be tuned for other applications in root and shoot biology, including trees and their stele and secondary cell wall architecture in both angiosperms and gymnosperms, anthers and other tissues undergoing secondary cell wall deposition. It may also have applications to mammalian systems including the human gut or other radially symmetrical structures.

Together, our results demonstrate that root anatomical barrier traits form an integrated and dynamic system shaped by interacting genetic and environmental factors. By enabling quantitative, cell-type-specific phenotyping across diverse experimental contexts, automated segmentation provides a scalable framework for systematically dissecting root barrier biology. This approach enables the integration of anatomical, genetic, environmental data to resolve how root systems adapt to complex conditions. More broadly, such frameworks will be essential for linking root structure to plant performance in dynamic environments.

## METHODS

### RADIX architecture

RADIX comprises a Vision Transformer encoder with self-supervised pretrained weights from Meta (DINOv3-S/16, HuggingFace checkpoint Facebook/dinov3-vits16-pretrain-lvd1689m)^25^ and a custom Dense Prediction Transformer (DPT) decoder following Ranftl et al.^30^. The encoder accepts a (3, 1024, 1024) input tensor and partitions the image into 16x16 non-overlapping patches, producing a 64x64 grid of 4096 patch tokens. Together with 1 [CLS] token and 4 register tokens (DINOv3’s standard configuration), this forms a sequence of 4101 tokens with embedding dimension 384. DINOv3 uses rotary position encoding (RoPE), which encodes each token’s spatial location through rotations applied inside the self-attention operation rather than as a fixed learned vector added to the input. This allows the encoder to run at 1024x1024 without re-interpolating positional embeddings from the pretraining resolution. The encoder consists of 12 transformer blocks, each with multi-head self-attention (6 heads) and an MLP sublayer.

The DPT decoder taps the patch-token output of layers 3, 6, 9, and 12 of the encoder, four evenly spaced layers spanning shallow to deep features. At each tap, the [CLS] and register tokens are discarded and the remaining 4096 patch tokens are reshaped to a (384, 64, 64) feature map. Each tap is projected through a 1x1 convolution to its target channel width (96, 192, 384, and 768 channels for the shallow-to-deep taps) and then resampled to its target stride. The shallowest tap is 4x upsampled with a transposed convolution to stride 4 (256x256), the second is 2x upsampled to stride 8 (128x128), the third is kept at the encoder’s native stride 16 (64x64), and the deepest is downsampled with a strided 3x3 convolution to stride 32 (32x32). A 3x3 convolution then projects all four maps to a unified width of 256 channels.

The four feature maps are progressively fused from deepest to shallowest through four fusion blocks, each consisting of residual convolutional units followed by 2x bilinear upsampling. This design ensures that the final per-pixel prediction is informed both by the precise local cellular boundaries visible in the shallow features and by the long-range anatomical context encoded in the deep features, while the skip connections at every scale stabilize learning by letting each decoder block refine the existing representation rather than reconstruct it from scratch. After four fusions, the decoder output is at stride 2 (512x512), which passes through a 256, 128, 32 channel reduction with two 3x3 convolutions, a 1x1 segmentation head producing 7 channels, and a final 2x bilinear upsampling to recover the input resolution. The result is a (7, 1024, 1024) class-logit tensor over the six anatomical classes (epidermis, exodermis, cortex, aerenchyma, endodermis, stele) plus a background class. Argmax across the class dimension produces the discrete anatomical class map.

### Image preprocessing

During model training and application, each sample consists of three single-channel TIFF images acquired through the DAPI, FITC, and TRITC fluorescence filter channels. The three channels are stacked into a single (H, W, 3) array, mapping TRITC, FITC, and DAPI to RGB. Each channel is independently normalized to [0, 1] by clipping to its 1st and 99.5th intensity percentiles and rescaling linearly; absent channels are set to a constant zero. Percentile normalization is performed per image rather than per dataset, so that intensity scaling is robust to the substantial differences in absolute brightness across imaging platforms. No background subtraction, deconvolution, or denoising is applied. Images are then resized so that the longer edge is 1024 pixels (preserving aspect ratio) and the shorter edge is zero-padded on the right or bottom to produce a 1024x1024 output; padding metadata is retained so that predictions can be cropped and rescaled back to native resolution for downstream analysis.

### Augmentation pipeline

Building on our large-scale dataset, data augmentation expands the effective training set even more by applying randomized, biologically plausible perturbations on the fly, forcing the model to learn features that generalize across staining quality, microscope settings, and sectioning artefacts, rather than memorizing the training images after hundreds of training epochs. Different augmentation was applied online for every epoch with the *Albumentations* and *Ultralytics* library during training, and the same pipeline was shared across all models compared in Figure 2 with proper random number generator seeding, so performance differences reflect architectural choices rather than augmentation tuning. No augmentation is applied during model application.

Geometric augmentations randomly altered the spatial layout of the image (see **Supplementary Fig. 3**). These were RandomRotate90 (p = 0.5), horizontal flip (p = 0.5), vertical flip (p = 0.5), an Affine transform combining translation (±10% of image size), scaling (0.7-1.3x), rotation (±45°), and shear (±10°) applied jointly (p = 0.7), and ElasticTransform (alpha = 120, sigma = 12; p = 0.3), which mimics the subtle distortions introduced during sectioning and mounting. Intensity augmentations randomly altered brightness and texture to mimic exposure, focus, and sensor differences across microscopes. These were RandomBrightnessContrast (brightness shift ±0.3, contrast shift ±0.3; p = 0.6), RandomGamma (γ ∈ [0.7, 1.5]; p = 0.3), GaussianBlur (kernel size 3-7 px; p = 0.2), and GaussNoise (per-pixel σ ∈ [0.01, 0.08] on the normalized image; p = 0.4). To promote robustness against single-channel staining failures and intensity differences between imaging platforms, ChannelDropout (exactly one input channel zeroed; p = 0.2) and ChannelShuffle (random permutation of the three channels; p = 0.2) were applied. After all augmentations, the image was resized to 1024x1024.

### Training procedure

RADIX was trained on a single NVIDIA H100 80 GB GPU using PyTorch 2 and PyTorch Lightning. We used the AdamW optimizer (weight decay 1 x 10^-4^) with two learning rates, since the encoder was pretrained and needed only gentle fine-tuning while the decoder was trained from scratch. The encoder learning rate was 1 x 10^-5^ and the decoder learning rate was 1 x 10^-4^, both following a cosine annealing schedule that smoothly decreased the learning rate to near zero over 200 epochs. Training stopped early if validation loss failed to improve for 15 consecutive epochs. Per-GPU batch size was 16. The random seed was fixed at 42.

The training objective summed four complementary losses; each applied to the full seven-class (six anatomical structure class and a background class) softmax output and weighted equally. Dice loss maximizes per-class region overlap and remains effective under class imbalance, which matters here because background pixels far outnumber small anatomical structures such as the endodermis. Focal loss redirects learning toward harder pixels by downweighting confidently classified ones, helping with fine structures such as the thin walls bounding aerenchyma lacunae. Cross-entropy loss (equal class weights) is the standard per-pixel classification objective. Lovász-softmax loss is a differentiable approximation of the intersection-over-union (IoU) metric and encourages sharper class boundaries. An ablation of Lovász-softmax loss term is provided in **Supplementary Table 7**.

### Comparison architectures

All seven comparison architectures were trained with the same data, loss combination, augmentation pipeline, optimizer, schedule, batch size, and epoch as RADIX. They differed in pre-training scale and the architectural choice.

The two convolutional baselines paired the segmentation_models_pytorch UNet++ decoder with ImageNet-pretrained ResNet34 (24.4 M parameters) and ResNet50 (32.5 M parameters) encoders. The three foundation-model variants probed which component of RADIX mattered most. DINOv2-DPT used the same DPT decoder as RADIX but swapped the DINOv3-S/16 encoder for DINOv2 (vit_small_patch14_dinov2 from timm, pretrained on LVD-142M), isolating the effect of the newer encoder. DINOv2-MS-Linear and SegDINO replaced the DPT decoder with simpler heads (a multi-scale linear head following the Meta MS-Linear recipe taps the last four transformer blocks, paired with DINOv2; and a small per-patch MLP head paired with DINOv3-S/16), isolating the effect of the decoder.

The MicroSAM+UNETR variant paired the SAM ViT-B encoder pretrained on a large microscopy corpus (∼89 M parameters)^36^ with a UNETR decoder^38^ tapping encoder blocks 3, 6, 9, and 12 and progressively upsampling them through ConvTranspose blocks with skip connections (∼9.6 M parameters). Because full fine-tuning of ViT-B at 1024x1024 exceeded GPU memory, we adapted the frozen encoder with LoRA adapters of rank 4 (α = 1.0) on every attention projection and trained only the LoRA parameters, the UNETR decoder, and the segmentation head (∼9.9 M trainable parameters), with the batch size reduced from 16 to 8.

The instance-segmentation baseline was YOLO26m-seg (Ultralytics)^39,52,53^, fine-tuned from a COCO-pretrained checkpoint to detect six object classes: the outer epidermis contour, aerenchyma, outer and inner endodermis contours, and outer and inner exodermis contours. Annotations were provided as filled polygons rather than per-pixel class maps. At inference, the predicted instance polygons were rasterized and combined by subtraction between nested contours to reconstruct pixel-level semantic masks over the same six anatomical classes used by the other seven models, enabling direct comparison on the same anatomical classes.

### Evaluation metrics

Throughout this work we report semantic segmentation accuracy as intersection-over-union (IoU), the standard metric in computer-vision benchmarks for dense pixel-wise prediction tasks^54–56^. For a predicted mask and its matched ground-truth mask, IoU is defined as:

IoU = TP / (TP + FP + FN),

where each pixel falls into one of three categories: a true positive (TP) is correctly predicted as belonging to the class, a false positive (FP) is predicted to belong to the class but does not in the ground truth, and a false negative (FN) is a ground-truth pixel that the model missed. IoU penalizes both over-prediction and under-prediction symmetrically. We chose IoU over precision (TP / (TP + FP)) and recall (TP / (TP + FN)) because each of those captures only one of the two failure modes.

We compute IoU independently for each of the six anatomical classes (epidermis, exodermis, endodermis, cortex, aerenchyma, and stele) and report their unweighted mean as the per- sample mean IoU (mIoU). The unweighted mean was chosen deliberately to prevent larger structures such as the cortex from dominating the overall score.

### Fluorescence intensity extraction and downstream trait quantification

Downstream trait quantification was performed directly on predicted segmentation masks without manual correction. For each cross-section, endodermis and exodermis tissue masks were derived by subtracting the inner boundary polygon from the outer boundary polygon (ring subtraction), yielding filled tissue-layer masks. Aerenchyma masks were used directly as predicted.

To isolate cell-wall fluorescence signal from cell lumen contributions, a user-defined intensity threshold was applied within each tissue mask on a per-batch basis. The threshold was selected to exclude low-intensity pixels while retaining cell-wall signal and was held constant across all samples within a given experimental batch. The upper intensity bound was fixed at the maximum value of the source bit depth (65,535 for 16-bit images; 255 for 8-bit images acquired on the Zeiss LSM 880 [out-of-distribution]), so no upper clipping was applied. Mean FITC and TRITC intensities above the threshold were computed per mask region and used as proxies for suberin and lignin deposition, respectively.

Aerenchyma ratio was computed as the total cross-sectional area of predicted aerenchyma polygons divided by the cross-sectional area of the whole root polygon for each section.

To validate that predicted mask boundaries introduce negligible error into downstream trait values, all four fluorescence intensity traits and aerenchyma ratio were extracted independently from both predicted masks and expert ground-truth annotations across all held-out test samples (n = 185 for fluorescence traits; n = 148 monocot samples for aerenchyma ratio). Agreement between predicted and ground-truth trait value was assessed by linear regression; the coefficient of determination (R²) and regression slope are reported.

### Plant material and growth conditions

Monocot species (millet, rice, and sorghum) were cultivated across multiple growth systems to introduce biologically relevant variation in root morphology and anatomical barrier formation. Growth systems included semi-hydroponic rhizotrons, rockwool-based rhizotrons, paper pouch systems, sand-based Falcon tube systems, and clay-based pot systems. Dicot species (tomato) were grown under sterile plate-based conditions (see below).

Semi-hydroponic rhizotrons consisted of chambers filled with 4 mm glass beads as an inert support matrix, with water supplied as the sole growth medium. Rockwool rhizotrons consisted of chambers in which plants were grown on rockwool as the supporting substrate. Paper pouch systems consisted of vertically oriented pouches supplied with water for seedling growth. Sand- based systems consisted of 50 mL Falcon tubes filled with sand and watered as needed. Clay pot systems consisted of greenhouse-grown plants cultivated in a clay-based substrate.

All cereal seeds were surface sterilized in 2.5% hypochlorite for 15 min with gentle agitation, rinsed five times with sterile deionized water, and treated overnight with 5% Captan fungicide, followed by an additional five washes. Seeds were then pre-germinated in a moist environment at 28 °C overnight prior to transfer into growth systems.

Across all growth systems, cereals were cultivated for 7 days under a 12 h light / 12 h dark photoperiod with a programmed diurnal cycle simulating sunrise, morning, noon, afternoon, and sunset, based on representative climate data from Senegal and Ethiopia. Maximum daytime temperature was 32 °C, and nighttime temperature was 19°C.

Seeds of *Oryza sativa* cv. Kitaake were imbibed in the dark for 2 days at 28°C and transferred to pots (15 cm diameter x 18 cm height) filled with Profile® Greens Grade™. Plants were grown in a greenhouse (UC Riverside) at 28°C for 16 h during the day and 25°C for 8 h at night. Twenty- one-day-old plants were subjected to water withholding for 7 days prior to sampling.

Tomato hairy root cultures were grown on media containing 4.3g /L Murashige and Skoog (MS) with Vitamins (Plant Cell Labs, MSP09-50LT) containing 0.5g/L MES buffer (Sigma, M8250- 250G),30g/L sucrose (Fisher Scientific, S5-500), and 10g/L agar (Difco, 214530), adjusted to a pH of 5.7 using 1M potassium hydroxide (Fisher Scientific, P250-500). Hairy roots were grown for 2-3 weeks at which point 1.5 cm roots tips were collected for downstream processing. All *Solanum* seeds were sterilized for 20 min in 50% (v/v) commercial bleach. Sterilized seeds were subsequently grown on media containing 4.3g/L MS with Basal Salts (Plant Cell Labs, MSP01- 50LT) and 0.5g/L MES buffer, adjusted to a pH of 5.7 using 1M potassium hydroxide. Hairy root cultures and all *Solanum* species seedlings were grown on their respective media in 12 x 12- cm^2^ petri dishes in a 23°C growth chamber. Seedlings were supplemented with 16h/8h light/dark photoperiod.

### Microbial inoculation

Sorghum plants grown in semi-hydroponic rhizotrons were cultivated either with or without microbial isolates. Microbial inoculation was performed at the time of seed transfer into growth systems (0 DAS). Bacterial cultures were grown in Tryptic Soy Broth (TSB) to an optical density at 600 nm (OD_600_) of 0.6, pelleted by centrifugation at 3,000 x g for 15 min, and resuspended in sterile water to minimize carry-over effects of the growth medium. A total volume of 1 mL of the washed inoculum was applied per semi-hydroponic rhizotron.

### Genetic materials and mutant generation

*WOX10* loss-of-function mutants were generated using CRISPR-Cas9 genome editing in rice (*Oryza sativa* cv. Kitaake) through *Agrobacterium*-mediated transformation. Guide RNA targets (guide1: CCCAGCTTGCAGTCGCAGTG; guide2: CAACAGCGGCATGGTGAACC) were designed within the *WOX10* coding sequence using CHOPCHOP selecting Kitaake v3.1 genome. The synthesized guide sequences were cloned into a Gateway entry vector under the control of the *OsU6.1* promoter and subsequently recombined into the plant expression vector pBY02-Cas9, which carries Cas9 and a hygromycin resistance gene. The final construct was introduced into *Agrobacterium tumefaciens* strain EHA105 and used to transform rice calli derived from mature embryos. Putative *wox10* mutants were screened by PCR amplification of the *WOX10* target region followed by Sanger sequencing. Homozygous Cas9-free mutant lines in the T2 generation were advanced and used for phenotypic characterization of root anatomical traits.

Tomato sterile hairy roots cultures were generated following the protocol from Ron et al. 2014^57^ Tomato cv. M82 wild type and *myb92-1* CRISPR-Cas9 knockout mutant lines were previously generated and obtained from Cantó-Pastor et al. 2024^5^. All other *Solanum* species were obtained from the Tomato Genetics Resource Center at UC Davis or the Lippman Lab at Cold Spring Harbor Laboratory (**Figure 1, Supplementary Data 1 and Supplementary Table 1**).

### Root sampling and sectioning

After growth, root segments were excised from defined regions along the root axis. For most experiments, sections were collected at defined distances from the root tip (e.g., 0-1 cm, 1-2 cm, 2-3 cm, up to ∼10-12 cm), capturing developmental gradients along the root. In addition, specific regions such as crown roots were sampled where applicable. Sampling ages ranged primarily from 5-7 days for seedling systems, with additional experiments conducted at later stages (e.g., 21 days for greenhouse-grown rice). Sample ages and regions are recorded in the dataset metadata (**Supplementary Data 2**). Root fragments were embedded in 4% (w/v) agarose and fixed in formaldehyde-acetic acid-alcohol (FAA) or 4% paraformaldehyde (PFA). Samples were rehydrated through a graded ethanol series (70%, 50%, 30%, and 10% ethanol; ≥30 min per step) and sectioned using a vibratome to obtain transverse root sections with a thickness of 150-200 µm. For inclusion in model training or testing whether the model could successfully distinguish a reduced-suberin phenotype in tomato roots, samples were collected from 8-to-10- day-old seedling primary roots within 1 cm of the hypocotyl junction.

### Histological staining

Sections were subjected to a sequential triple-staining protocol adapted from Sexauer et al. ^16^. Samples were stained with Basic Fuchsin for 15 min, followed by two rinses in ClearSee, with the second rinse lasting at least 30 min. Sections were then briefly rinsed once with deionized water and stained with Fluorol Yellow for 15 min, followed by a quick rinse in deionized water. Sections were subsequently stained with Calcofluor White for 15 min, followed by a brief wash in 50% ethanol and two final rinses in deionized water. After staining, sections were stored in water or 50% glycerol until imaging.

To demonstrate model performance in dicots for cell wall analysis, sections sampled from within one centimeter of the hypocotyl junction in wild type M82 and *slmyb92-1* seedlings were stained for suberin following the protocol from Cantó-Pastor et al.^5^. Briefly, root sections were cleared with methanol and subsequently stained with Calcofluor White (0.1% w/v in methanol) for 10 min and rinsed once with methanol. The sections were then incubated with Fluorol Yellow 088 (0.05% w/v in methanol) for one hour, then washed twice more with methanol. The washed sections were lastly counterstained with Aniline Blue (0.5% w/v in methanol) for 30 min. All staining steps were performed at room temperature in the dark.

### Fluorescence imaging

Stained root cross-sections were imaged on three fluorescence microscopy platforms. The majority of images (n = 1,501) were acquired on an Olympus IX83 inverted microscope equipped with a Cicero spinning-disk confocal unit (50 um pinhole diameter, 250 um disk pitch) and a Hamamatsu ORCA Flash LT3 sCMOS camera (6.5 um pixel pitch; effective pixel size 0.325 um at 20x), using a 20x air objective (NA 0.80). Fluorescence was excited with 405, 470 and 555 nm laser lines and detected through bandpass emission filters matched to each dye (DAPI channel for Calcofluor White, FITC for Fluorol Yellow 088, TRITC for Basic Fuchsin). Image acquisition and z-stack control were performed with cellSens Dimension 4.3. Z-stacks were collected per section with the number of planes and axial step size adjusted per experimenter; acquisition parameters (exposure times, laser power, z-step) were held constant within each imaging session and are reported in **Supplementary Data 2**. An additional 159 images were acquired on a BioTek Cytation C10 confocal imaging reader using a 20x air objective (NA 0.450; effective pixel size 0.325 um), with DAPI, GFP, and TRITC fluorescence cubes corresponding to Calcofluor White, Fluorol Yellow 088, and Basic Fuchsin, respectively, controlled with Gen5 v3.16 software. A further 35 images were acquired on a Zeiss LSM 880 laser scanning confocal microscope using a 20x air objective (effective pixel size 0.461 um; 1024x1024 pixels), with 405, 488, and 561 nm laser lines. Fluorescence was collected through a pinhole set to 1.0 Airy Unit using a photomultiplier tube (PMT) detector with emission bandwidths of 410-500 nm (DAPI), 493-541 nm (FITC), and 566-612 nm (TRITC); acquisitions used scan speed 9 (3.77 s per frame). Image acquisition was controlled with ZEN Blue software.

For Olympus IX83/Cicero images, z-stacks were processed using extended focal imaging (EFI) to generate a single in-focus composite per section, followed by deconvolution to improve signal clarity. No manual contrast adjustment or other subjective image manipulation was applied after EFI and deconvolution. Individual fluorescence channels (DAPI, FITC, TRITC) were exported as separate lossless grayscale image files as input for model training and inference.

### Dataset annotation

Root cross-section images used for model training and evaluation were obtained from multiple cereal species, including sorghum, rice, and pearl millet, encompassing multiple genotypes per species. Primary ground truth annotations were generated by an external professional image annotation service, Labellerr^29^. All annotations were subsequently reviewed and quality-controlled by the authors with domain-specific training to ensure biological plausibility and internal consistency.

The externally generated and internally reviewed annotations were used for supervised model training and for quantitative performance evaluation. In addition, a small independent subset of images (approximately 10-20) was manually annotated by authors to provide expert biological validation of model predictions. This expert-annotated subset was not used for model training and served solely for qualitative assessment and confirmation of biological interpretability.

### Statistics and Reproducibility

For the sorghum genotype comparison (**Fig. 5a**), differences among the 10 selected genotypes were assessed by one-way ANOVA followed by Tukey’s honestly significant difference (HSD) post-hoc test, with compact letter displays (CLD) generated at α = 0.05. Fluorescence intensities are shown normalized to the mean of genotype 23 (Teshale) to facilitate visual comparison; statistical analysis was performed on raw (unnormalized) values.

For microbial inoculation experiments (**Fig. 5b** and **Supplementary Fig. 7**), each microbial treatment was extracted from a larger experimental series alongside shared control samples spanning the matching experiments. To account for experiment-to-experiment variability while correctly isolating within-experiment treatment effects, a mixed-effects linear model was fitted by restricted maximum likelihood (REML) to raw thresholded fluorescence intensities with Treatment as a fixed effect and Experiment as a random effect (value ∼ C(Treatment) + (1|Experiment),REML). Pairwise comparisons were made between each treatment and the Control only; treatment-to-treatment comparisons were not performed as these comparisons are confounded with the experimental batch. P-values were corrected for multiple comparisons across all traits and treatments using the Benjamini-Hochberg false discovery rate (FDR) procedure.

For the Wox10 experiment (**Fig. 5c**), fluorescence intensities and aerenchyma ratios were first averaged across cross-sections within each plant to obtain one biological replicate per plant (n = 3-5 plants per genotype per system). Differences among genotypes within each growth system were assessed by one-way ANOVA. Post-hoc comparisons against the WT control were performed using Dunnett’s test for aerenchyma ratio.

For the tomato *myb92-1* experiment (**Fig. 5d**), a two-sided Mann-Whitney U test was used to compare exodermal suberin (FITC) fluorescence intensity between Control (n = 4) and *myb92- 1* (n = 6) plants, as data could not be assumed to be normally distributed given the small sample size.

## Acknowledgements

Work in this project was funded by the Gates Foundation (Seattle, WA, USA) via grant OPP1082853 “RSM Systems Biology for Sorghum.” to NIOO and “PROMISE II: Promoting microbes for integrated Striga eradication” (INV-047255); and NSF-IOS 2118017 and NSF-PGRP-IOS-1856749 to JBS and SMB. We also acknowledge support from NIH R21CA313790. We thank Zachary Lippman for Solanum seed provision; Hana Lubin, Alex Kim-Fonkalsrud, and Nisha Ganesan for assistance in preparing sections and imaging; Justin Leong, Rithika Oumapathy, Qiudi Cheng, Lola Stevens, and Jenni Pimentel for image annotations; and Taye Tessema for provision of sorghum seeds.

## Competing interests

The authors declare no competing interests.

## Data availability

The RADIX benchmark dataset generated in this study, comprising 1,695 fluorescence microscopy images of root cross-sections (DAPI, FITC, and TRITC channels) and their corresponding expert-curated six-class anatomical annotations, together with the training, validation, in-distribution test, and out-of-distribution (Zeiss) test splits and per-sample metadata, has been deposited at Zenodo under accession: DOI: 10.5281/zenodo.21347957 and is publicly available. All code for data preprocessing, model training, inference, and downstream trait quantification, including the RADIX architecture and the seven benchmarked comparison models, is also deposited at the same Zenodo data source and maintained at https://github.com/y222gu/radix. The repository includes the training configurations, augmentation pipeline, evaluation scripts, and trained model weights required to reproduce the results reported here. A condensed annotation protocol is provided as **Supplementary Information**. The DINOv3 pretrained encoder checkpoint used for initialization is publicly available from Meta (Hugging Face: facebook/dinov3-vits16-pretrain-lvd1689m). Plant materials, including the CRISPR-Cas9 *wox10* lines, are available from the corresponding authors upon reasonable request; the *slmyb92-1* line was obtained from Cantó-Pastor et al. ^5^ and other *Solanum* accessions from the Tomato Genetics Resource Center (UC Davis) and the Lippman Laboratory (Cold Spring Harbor Laboratory).

## Author contributions

Yifei Gu - Conceptualization, Investigation, Methodology, Software, Data curation, Formal analysis, Validation, Visualization, Project administration, Writing - original draft, Writing - review & editing

Stefan Sanow - Conceptualization, Investigation, Methodology, Data curation, Formal analysis, Validation, Visualization, Project administration, Writing - original draft, Writing - review & editing

Tamera Taylor - Conceptualization, Investigation, Methodology, Data curation, Formal analysis, Validation, Visualization, Project administration, Writing - original draft, Writing - review & editing

Kevin Morimoto - Conceptualization, Investigation, Methodology, Data curation, Validation, Visualization, Writing - review & editing

Adele Nemer - Investigation, Data curation, Validation, Writing - review & editing Dustin J Hadley - Investigation, Writing - review & editing

Syed Adeel Zafar - Investigation, Data curation, Resources, Writing - review & editing Lucas Demello - Investigation

Julia Bailey-Serres - Resources, Supervision

Randy P. Carney - Conceptualization, Funding acquisition, Project administration, Supervision, Writing - original draft, Writing - review & editing

Siobhán M. Brady - Conceptualization, Funding acquisition, Project administration, Supervision, Writing - original draft, Writing - review & editing

## Supplementary Figures

**Supplementary Figure 1.**
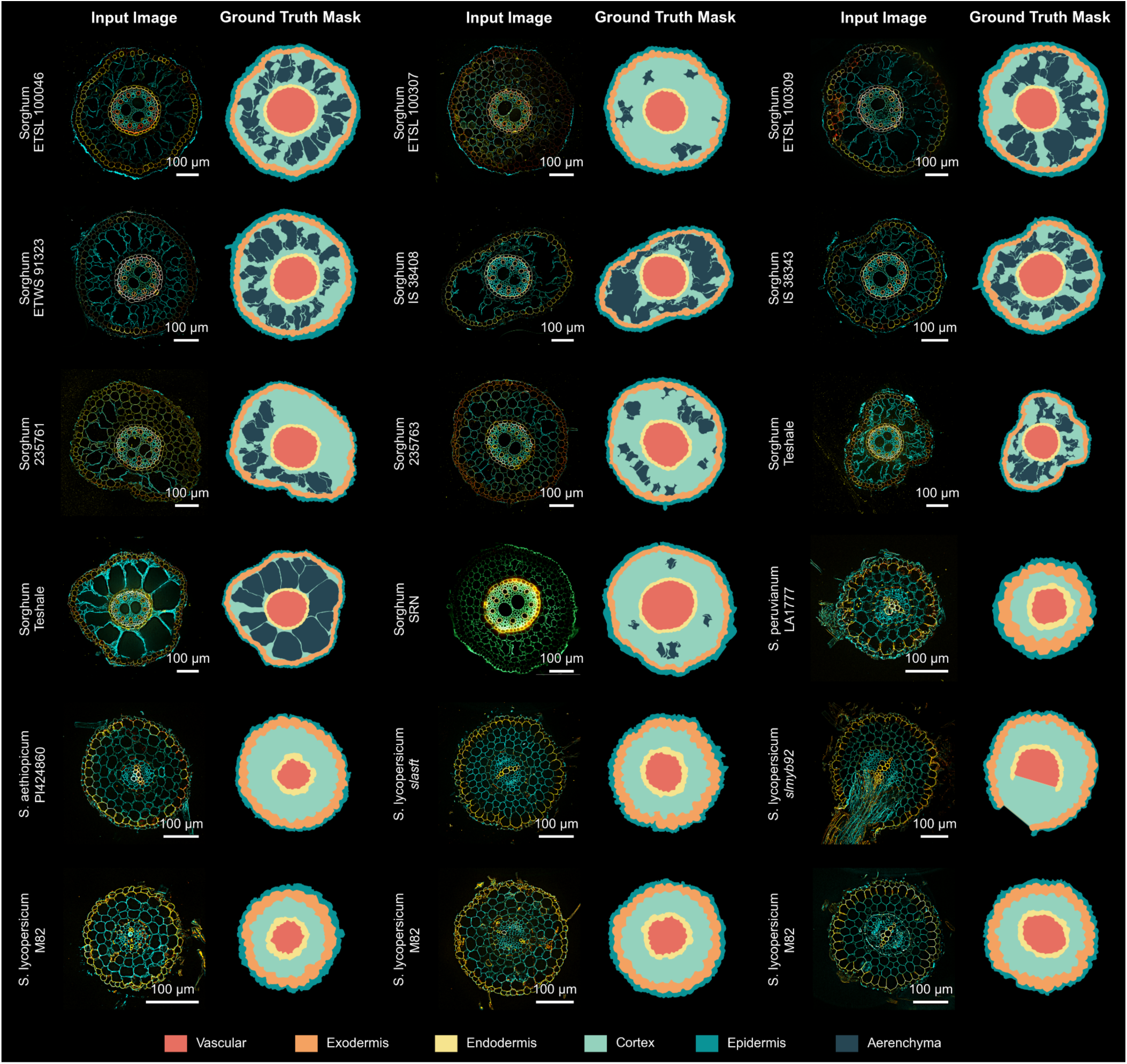
Additional representative samples in the training data set in three channel composites (DAPI in cyan, FITC in yellow, and TRITC in red) with corresponding ground truth annotation used during supervised training.

**Supplementary Figure 2.**
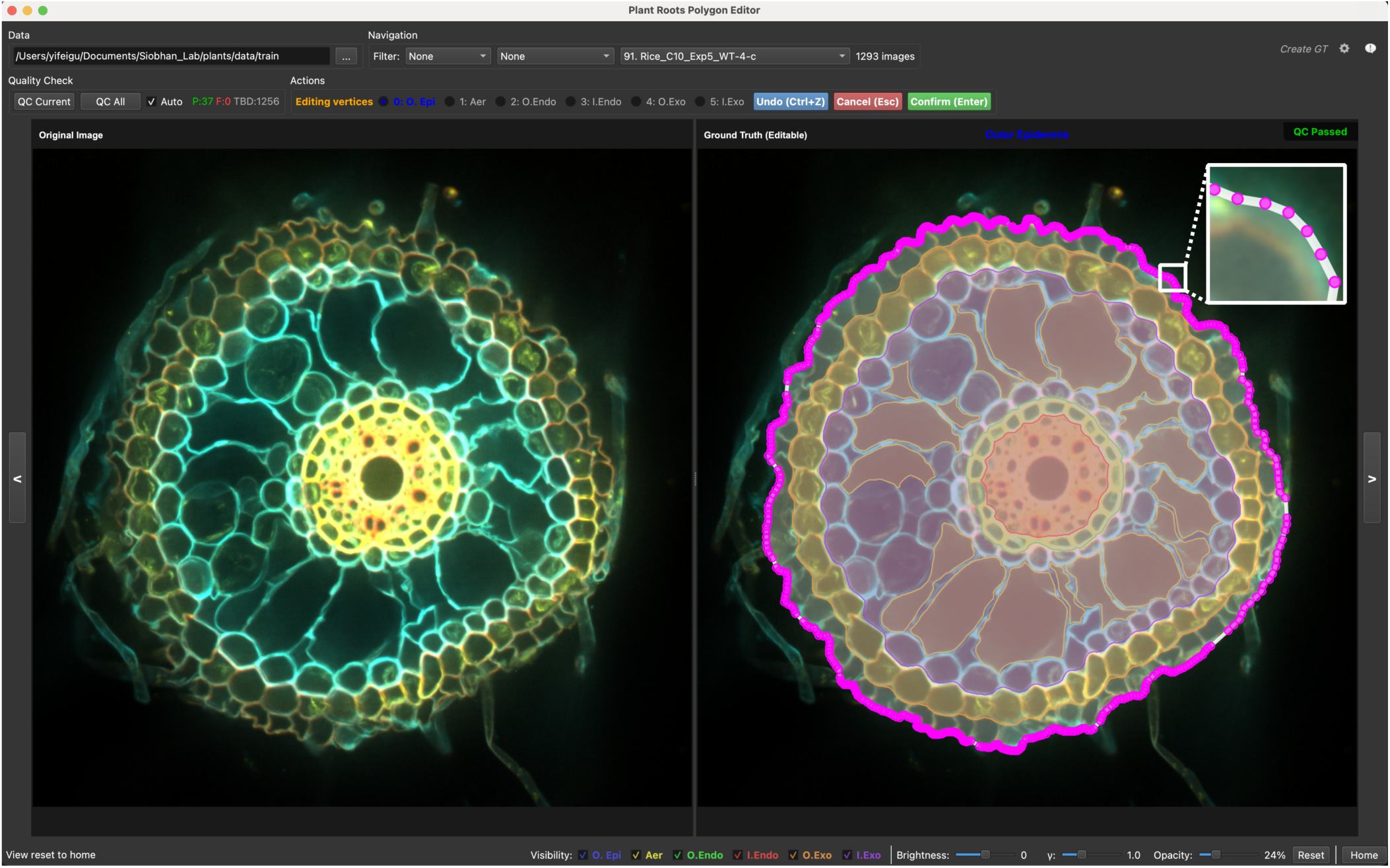
Annotation review tool. *A screenshot of the GUI built for rapid annotation review*.

**Supplementary Figure 3.**
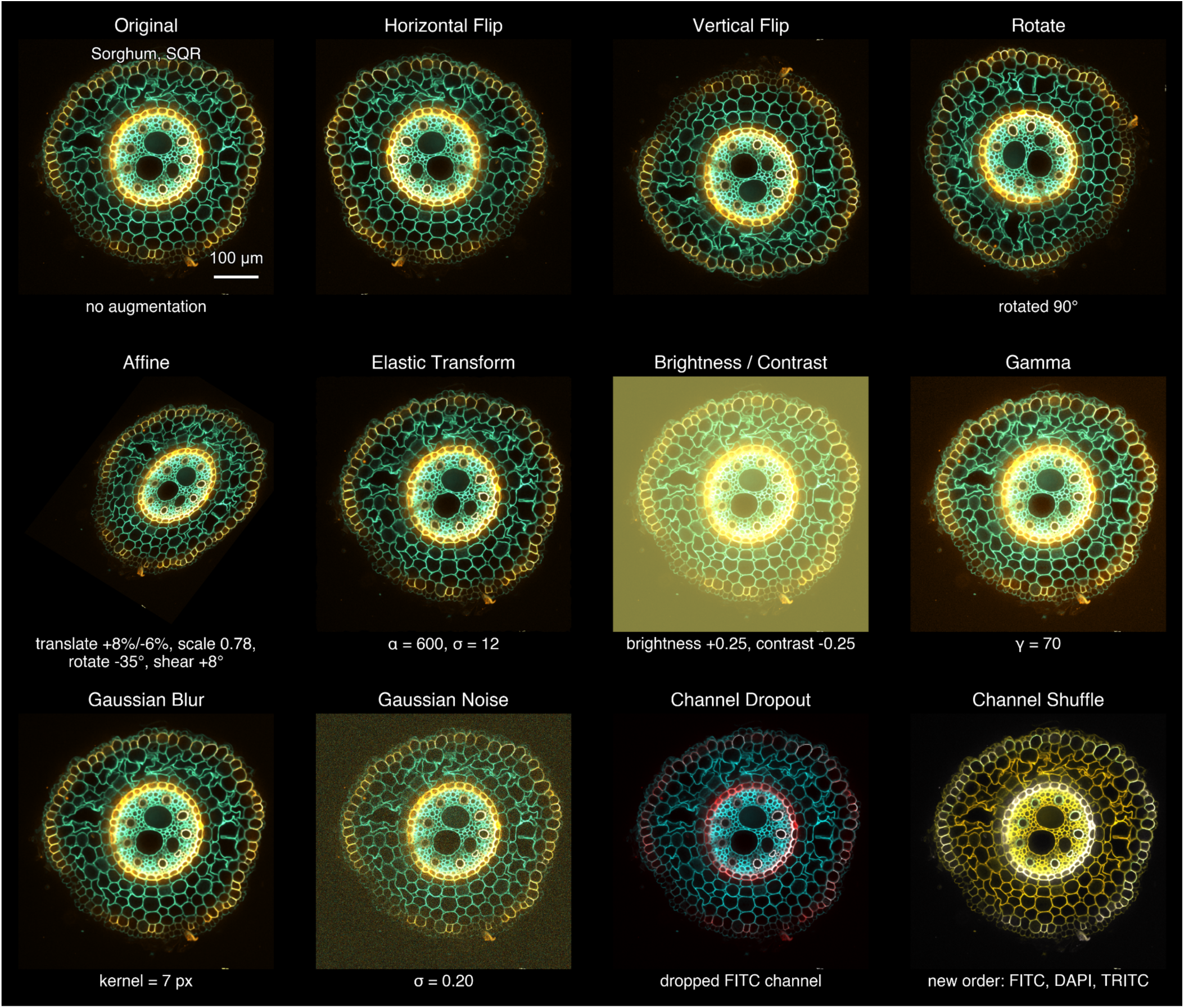
An example image is shown with different types of augmentation operation used during training. *Multiple of these operations could be applied with random parameters described in Methods during training. The top-left panel is the unaltered sample for reference; the remaining eleven panels show, in order, 90° random rotation, horizontal flip, vertical flip, affine transformation (translation, isotropic scaling, in-plane rotation, and shear), elastic deformation, brightness and contrast jitter, gamma adjustment, Gaussian blur, additive Gaussian noise, channel dropout (one of the three fluorescence channels set to zero), and channel shuffle (the three channels randomly permuted). The exact parameter values used in this figure are annotated below the corresponding image. All channels are displayed using the paper convention: DAPI in cyan, FITC in yellow, and TRITC in red, additively blended. Geometric operations (90° rotation, flips, affine, elastic) are applied identically to the image and to the semantic segmentation mask during training; Intensity and channel operations modify only the image. During training, each operation is sampled independently at the probabilities reported in the Methods section*.

**Supplementary Figure 4.**
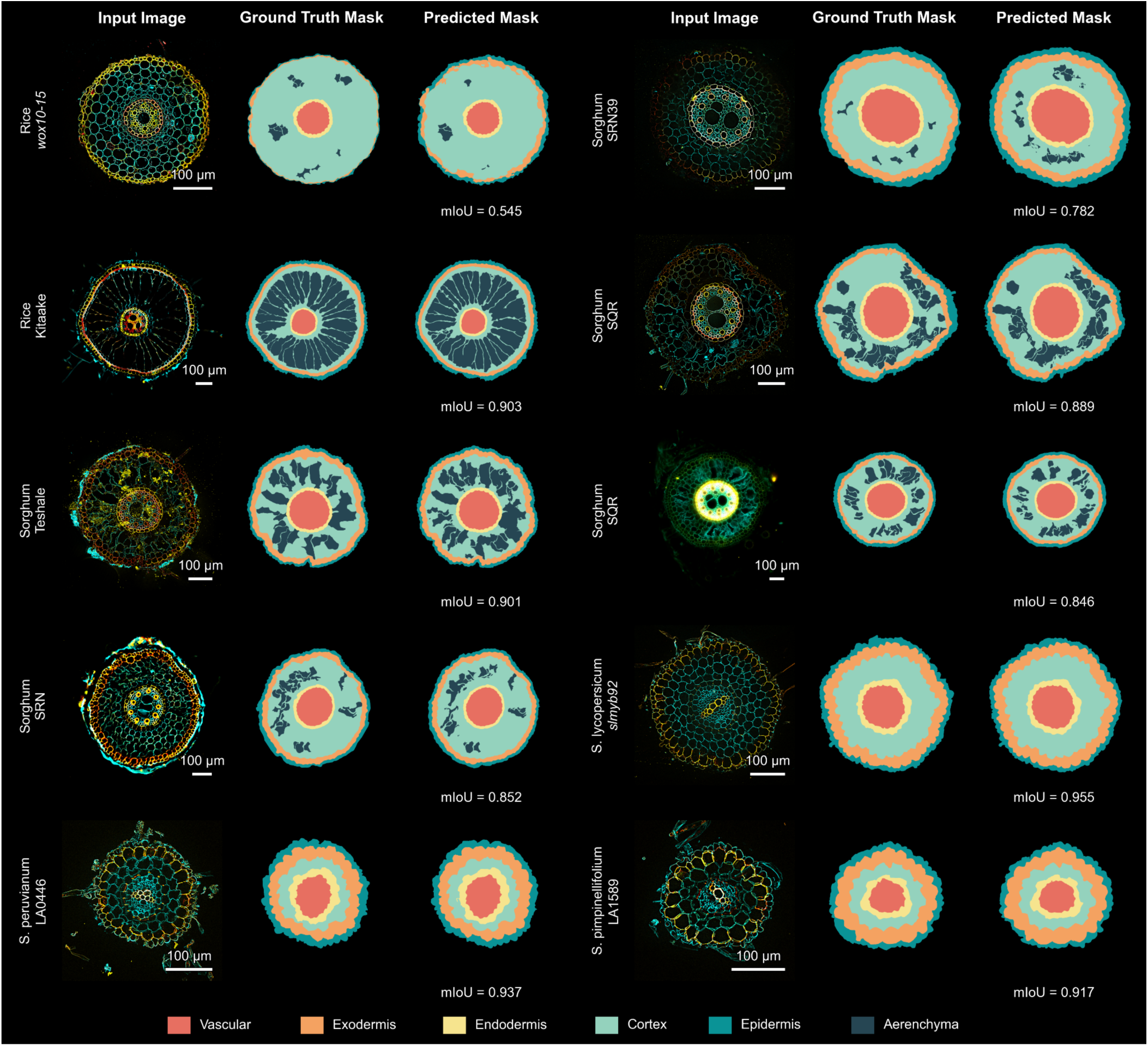
**Additional representative examples of model predicted mask vs. ground truth mask from test data sets.**

**Supplementary Figure 5.**
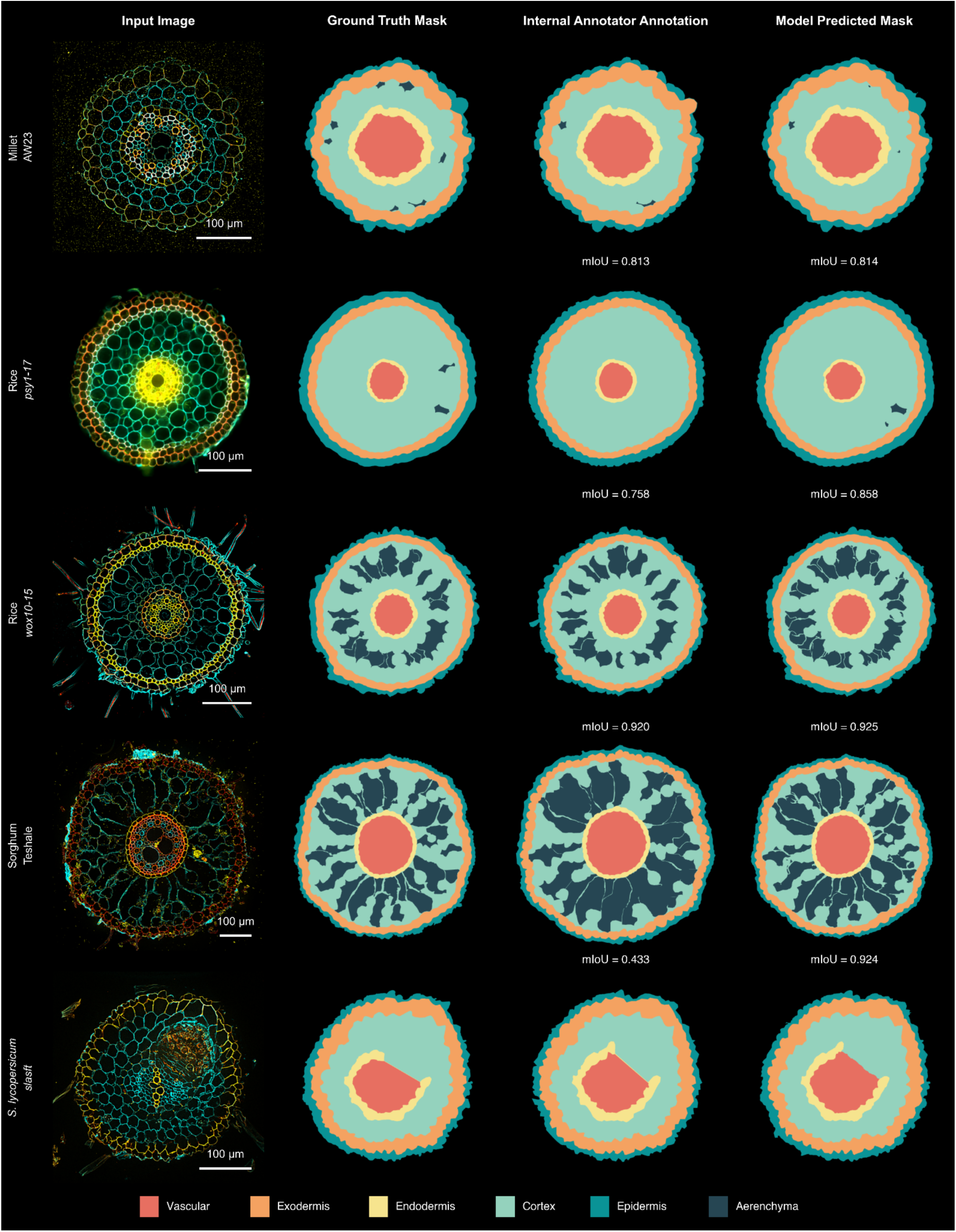
Visual comparison of ground-truth, internal-annotator annotations, and RADIX predictions on five representative test-set samples. *Each row shows one root cross-section. Column from left to right: (1) the input fluorescence composite (DAPI in cyan, FITC in yellow, TRITC in red); (2) the consensus ground-truth annotation used during training; (3) an independent re-annotation by a second expert plant biologist, blinded to the consensus ground truth, drawn from the balanced subset of sixteen test-set samples to estimate inter-annotator agreement* (Fig. 2d); *(4) the prediction of RADIX. Masks are colored by anatomical class using the palette shown in the bottom legend (epidermis, exodermis, cortex, aerenchyma, endodermis, stele). The five samples span both monocots (millet, rice, sorghum) and a dicot (Solanum lycopersicum), and illustrate the typical pattern observed across all sixteen re-annotated samples(**Sup. Table 5**)*.

**Supplementary Figure 6.**
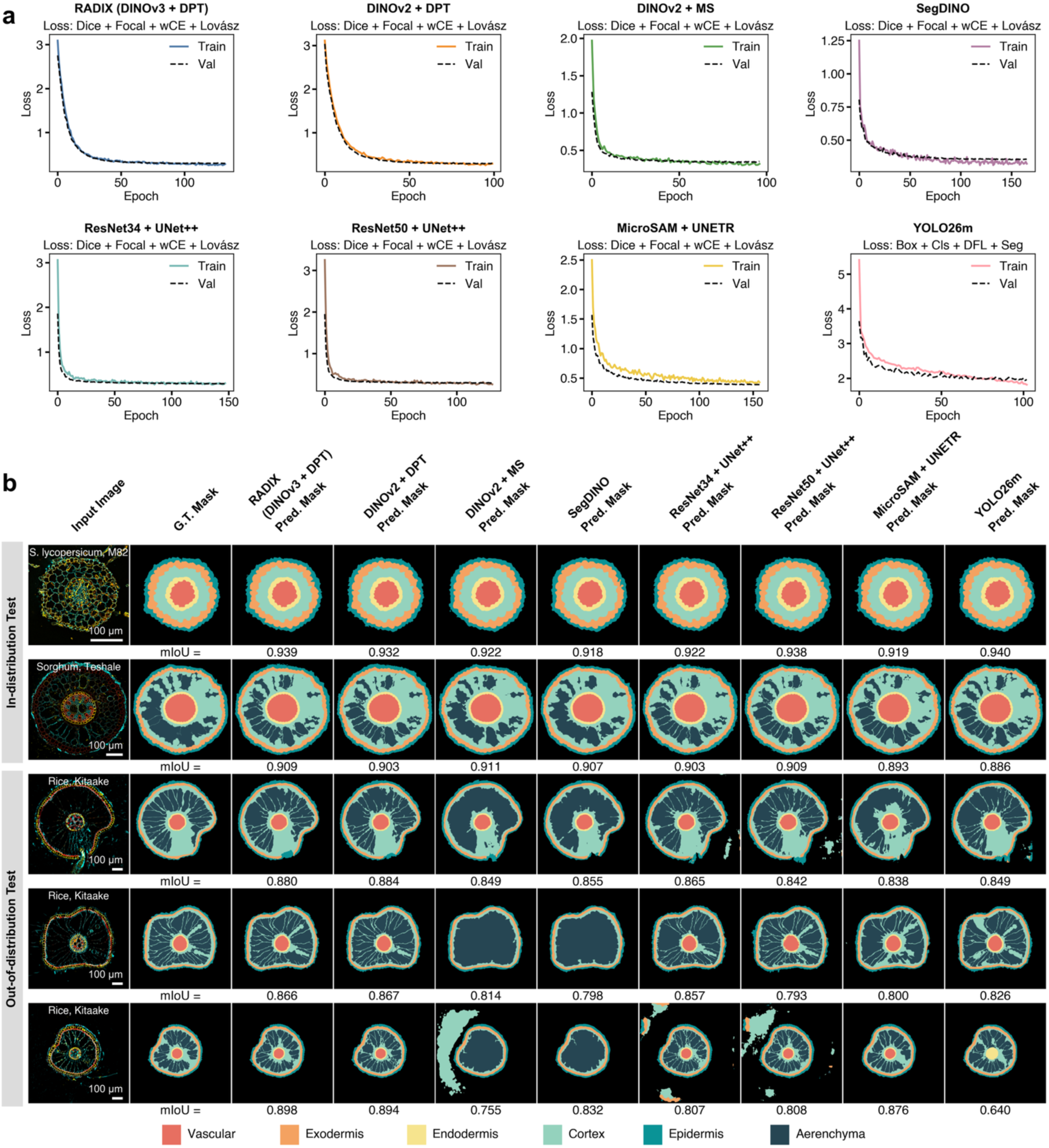
Architecture comparison of eight segmentation models. *(a) Training and validation loss curves for each model, with the loss formulation it was trained on annotated in the subplot title. All seven semantic models (RADIX = DINOv3-S/16 + DPT decoder; ResNet34 + UNet++; ResNet50 + UNet++; DINOv2-S/14 with the same DPT decoder; DINOv2 with a multi-scale linear head; SegDINO; MicroSAM ViT-B + UNETR) optimize the same Dice + Focal + cross-entropy (wCE) + Lovász combination at 1024x1024 resolution. YOLO26m-seg, an instance-segmentation model, is trained with the Ultralytics box + class + DFL + segmentation loss. Curves are recorded per epoch from the corresponding training log; training loss is shown in color and validation loss as a dashed black line. All eight models reach a low, stable plateau within the 200-epoch training budget (early stopping triggered when validation loss failed to improve for 15 consecutive epochs), indicating that the chosen budget is sufficient for convergence and that the comparison in panel b is not confounded by under-training. (b) Per-sample predictions on five examples: two in-distribution test images and three out-of-distribution Zeiss images that no model saw during training or validation. Columns from left to right show the three channels composite (DAPI in cyan, FITC in yellow, TRITC in red), the ground-truth mask, and the prediction of each model painted in the anatomy palette shown in the legend. The mean IoU printed below each prediction is averaged across the six anatomical classes. RADIX matches or exceeds every alternative on the two in-distribution samples and is the most robust model under the Zeiss out-of-distribution shift, while other architectures show characteristic failure modes there. DINOv2-MS-Linear incorrectly predicts exodermis on the Solanum M82 sample, ResNet50 + UNet++ produces a fragmented epidermis on one of the Zeiss rice samples, and YOLO26m-seg collapses the central root region on the Solanum M82 sample*.

**Supplementary Figure 7.**
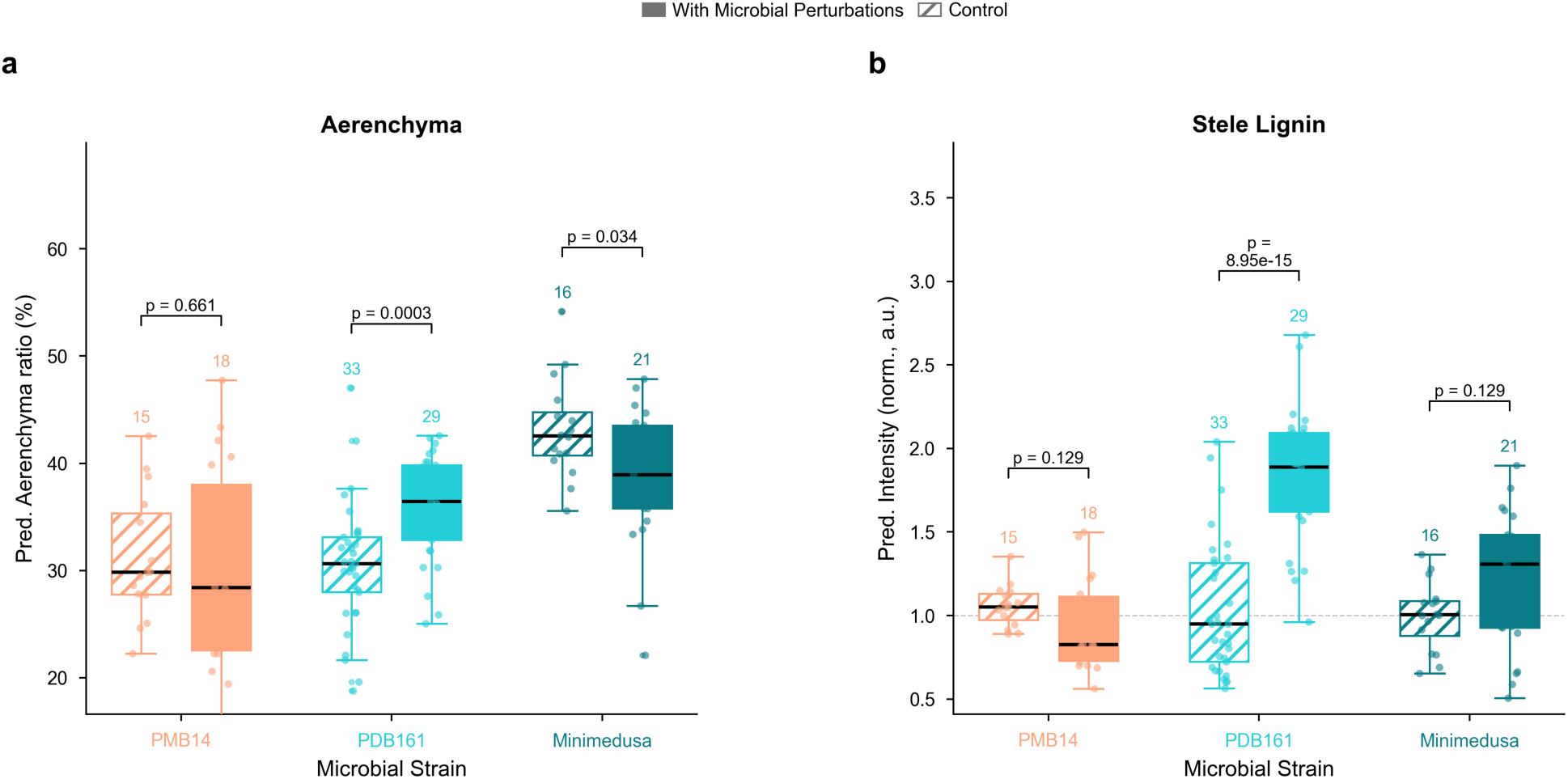
*Effects of microbial inoculants on sorghum root barrier traits: aerenchyma and stele lignin. Boxplots showing aerenchyma and stele lignin measurements in 7-day-old sorghum roots treated with three microbial isolates featured in* Figure 5b*, each compared to within-experiment controls. a, Aerenchyma ratio (cross-sectional area of air spaces relative to total root area, %). b, Stele (vascular) TRITC intensity (lignin proxy), normalized to the per-experiment control mean (dashed line at 1.0). These two traits are not shown for these isolates in* Figure 5b *and are reported here separately. Isolates: PMB14 (salmon, Exp. 1), PDB161 (cyan, Exp. 4), Minimedusa (teal, Exp. 9). Within-experiment controls are shown as hatched boxes to the left of each treatment group. Numbers above boxes indicate total cross-sections per group. Boxes span the interquartile range (IQR); center lines indicate medians; whiskers extend to the most extreme value within 1.5xIQR; individual data points are overlaid. P-values shown above brackets are BH-adjusted; statistical analysis follows the same approach as for* Figure 5b *(linear mixed-effects model with Experiment as random effect, Benjamini-Hochberg FDR correction across all traits and treatments), as described in detail in the **Methods***.

**Supplementary Figure 8.**
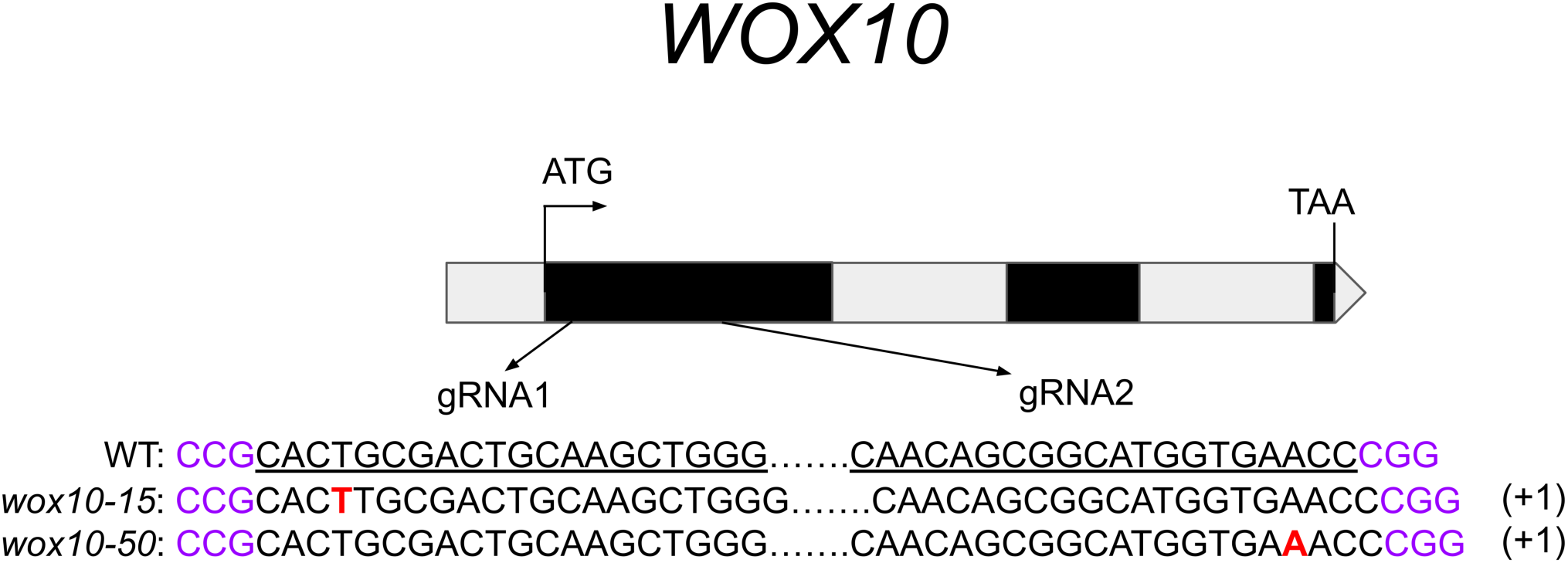
CRISPR-Cas9-mediated disruption of *WOX10* in *Oryza sativa*. *Schematic of the WOX10 (Os08g0242400) gene structure showing exons (black boxes) and introns (white boxes). ATG and TAA indicate the start and stop codons, respectively. Arrows indicate the positions of two guide RNAs (gRNA1 and gRNA2). Underlined sequences show the gRNA target sites; purple indicates the PAM sequence (CGG). The sequences of the wild-type (WT) and two independent CRISPR-Cas9 wox10 mutant alleles (wox10-15 and wox10-50) are shown below. Red nucleotides indicate the 1-bp insertions (+1) predicted to cause a frameshift mutation in each allele*.

## Supplementary Tables

**Supplementary Table 1.** Test sets diversity by species and genotype (n= 185 in- distribution, 35 out-of-distribution).

| Species | Genotype | In-distribution<br>Test | Out-of-distribution<br>Test | Total |
| --- | --- | --- | --- | --- |
| Millet | AW23 | 32 | 0 | 32 |
| Rice | Kitaake | 9 | 35 | 44 |
|  | *psy1-17* | 3 | 0 | 3 |
|  | *psy1-9* | 3 | 0 | 3 |
|  | *wox10-15* | 42 | 0 | 42 |
| Sorghum | SQR | 8 | 0 | 8 |
|  | SRN | 5 | 0 | 5 |
|  | SRN39 | 4 | 0 | 4 |
|  | Teshale | 42 | 0 | 42 |
| *Solanum lycopersicum* | M82 | 22 | 0 | 22 |
|  | *slasft* | 4 | 0 | 4 |
|  | *slmyb92* | 5 | 0 | 5 |
| *Solanum abutiloides* | Baker_Creek | 1 | 0 | 1 |
| *Solanum candidum* | 994750097 | 1 | 0 | 1 |
| *Solanum cheesmaniae* | LA1407 | 1 | 0 | 1 |
| *Solanum insanum* | INS1 | 1 | 0 | 1 |
| *Solanum peruvianum* | LA0446 | 1 | 0 | 1 |
| *Solanum pimpinellifolium* | LA1589 | 1 | 0 | 1 |

**Supplementary Table 2.** Per-class IoU of RADIX on the in-distribution and out-of- distribution test sets.

| Anatomical Class | In-distribution Test IoU<br>(mean ± s.d., n = 185) | Out-of-distribution Test IoU<br>(mean ± s.d., n = 35) |
| --- | --- | --- |
| Vascular | 0.979 ± 0.016 | 0.974 ± 0.009 |
| Exodermis | 0.878 ± 0.187 | 0.847 ± 0.084 |
| Endodermis | 0.912 ± 0.045 | 0.858 ± 0.046 |
| Cortex | 0.971 ± 0.040 | 0.982 ± 0.009 |
| Epidermis | 0.820 ± 0.131 | 0.738 ± 0.102 |
| Aerenchyma | 0.601 ± 0.265 | 0.835 ± 0.085 |

Note: Per-anatomical-class IoU (mean ± s.d. across samples) for RADIX on the in-distribution test set (n = 185 samples) and on the out-of-distribution set (n = 35 samples). Values correspond to Figure 2b and 2c. Aerenchyma is computed only over samples in which the class is present; Solanums lack the class entirely.

**Supplementary Table 3.** Per-class IoU and sample-level mIoU of RADIX, partitioned by species and by microscope.

| Group | n | Vascular | Exodermis | Endodermis | Cortex | Epidermis | Aerenchyma | mIoU |
| --- | --- | --- | --- | --- | --- | --- | --- | --- |
| *By species* |  |  |  |  |  |  |  |  |
| Rice | 92 | 0.973 ± 0.016 | 0.809 ± 0.241 | 0.874 ± 0.057 | 0.970 ± 0.045 | 0.742 ± 0.175 | 0.755 ± 0.179 | 0.854 ± 0.085 |
| Millet | 32 | 0.984 ± 0.006 | 0.930 ± 0.034 | 0.937 ± 0.021 | 0.972 ± 0.013 | 0.874 ± 0.042 | 0.286 ± 0.213 | 0.834 ± 0.045 |
| Sorghum | 59 | 0.987 ± 0.005 | 0.895 ± 0.104 | 0.920 ± 0.028 | 0.980 ± 0.041 | 0.835 ± 0.047 | 0.671 ± 0.206 | 0.883 ± 0.048 |
| Solanums | 37 | 0.974 ± 0.022 | 0.949 ± 0.028 | 0.922 ± 0.024 | 0.972 ± 0.018 | 0.864 ± 0.039 | n/a | 0.936 ± 0.018 |
| *By microscope* |  |  |  |  |  |  |  |  |
| Olympus | 161 | 0.980 ± 0.017 | 0.870 ± 0.199 | 0.916 ± 0.044 | 0.970 ± 0.043 | 0.814 ± 0.138 | 0.593 ± 0.271 | 0.868 ± 0.078 |
| C10 | 24 | 0.974 ± 0.010 | 0.930 ± 0.040 | 0.887 ± 0.039 | 0.983 ± 0.007 | 0.862 ± 0.046 | 0.675 ± 0.180 | 0.905 ± 0.040 |
| Zeiss | 35 | 0.974 ± 0.009 | 0.847 ± 0.084 | 0.858 ± 0.046 | 0.982 ± 0.009 | 0.738 ± 0.102 | 0.835 ± 0.085 | 0.872 ± 0.042 |

Note: Per-anatomical-class IoU and per-sample mIoU (mean ± s.d. across samples) for RADIX, broken down by species (top block) and by microscope (bottom block). Sample-level mIoU is the unweighted mean over the six anatomical structures for each sample. Rice pools both test sets (in-distribution and Zeiss out-of-distribution); the other species use the in-distribution test set only. Microscope rows are non-overlapping by construction; Zeiss is the held-out out-of-distribution row. n is the number of samples per row. Aerenchyma is computed only over samples in which the class is present; Solanums lack the class entirely. Values correspond to Figure 2d and 2e.

**Supplementary Table 4.**
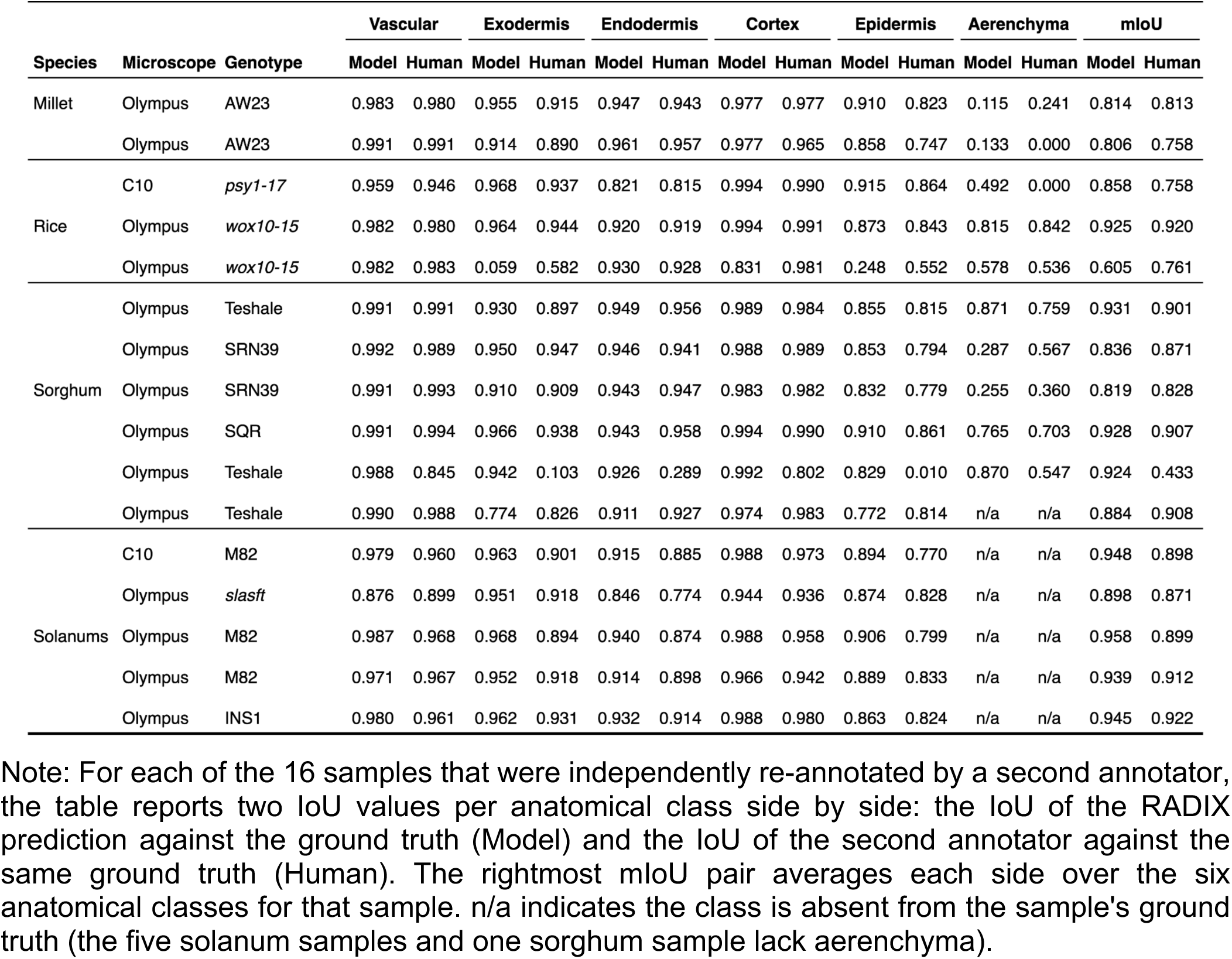
Inter-annotator IoU compared with RADIX prediction IoU, for each of 16 re-annotated samples.

**Supplementary Table 5.** Sample-level mIoU on the in-distribution and out-of-distribution test sets for the eight benchmarked architectures.

| Model | Encoder | Pretrained Dataset | In-distribution Test mIoU<br>(mean ± s.d., n = 185) | Out-of-distribution Test mIoU<br>(mean ± s.d., n = 35) |
| --- | --- | --- | --- | --- |
| RADIX | DINOv3-S/16 | LVD-1.69B | 0.873 ± 0.076 | 0.872 ± 0.042 |
| DINOv2 + DPT | DINOv2-S/14 | LVD-142M | 0.869 ± 0.071 | 0.872 ± 0.034 |
| ResNet50 + UNet++ | ResNet50 | ImageNet-1k | 0.868 ± 0.075 | 0.832 ± 0.055 |
| ResNet34 + UNet++ | ResNet34 | ImageNet-1k | 0.867 ± 0.078 | 0.854 ± 0.048 |
| YOLO26m-seg | YOLO26m | COCO | 0.866 ± 0.079 | 0.785 ± 0.112 |
| DINOv2 + MS-Linear | DINOv2-S/14 | LVD-142M | 0.858 ± 0.077 | 0.846 ± 0.036 |
| SegDINO-MLP | DINOv3-S/16 | LVD-1.69B | 0.856 ± 0.071 | 0.836 ± 0.042 |
| MicroSAM + UNETR | SAM ViT-B | SA-1B + LM | 0.827 ± 0.088 | 0.811 ± 0.059 |

Note: Sample-level mIoU (mean ± s.d. across samples) for every model in the Figure 2f, evaluated on the in-distribution test set (n = 185) and the Zeiss out-of-distribution set (n = 35). Sample-level mIoU is the unweighted mean over the six anatomical classes for each sample. The Encoder and Pretrained Dataset columns identify the encoder backbone and the dataset its weights were pretrained on. RADIX shows the highest IoU score on both in-distribution and out- of-distribution test sets.

**Supplementary Table 6.** Ethiopian Sorghum germplasm.

| ID | Genotype | Variety | Region/Origin |
| --- | --- | --- | --- |
| 1 | Wetetbegunchie | Local land race | Wollo |
| 11 | Bobered | Local Land Race | Benishangoul Gumuz |
| 17 | Framida | Striga-resistant | Purdue University |
| 23 | Teshale | Preferred Ethiopian Farmer Variety | Melkassa Agricultural Research Center |
| 37 | Jigurti | High Striga Germination Stimulant/Land Race | Jigurti |
| 42 | ETWS 91323 | Drought Tolerant, Low Germination Stimulant | Mondi/Menesibu |
| 45 | IS 38343 | Drought Tolerant | Oromia/Amhara |
| 50 | ETSL 100726 | Drought Tolerant | Melkassa Agricultural Research Center |
| 51 | ETSL 100576 | Drought Tolerant | Melkassa Agricultural Research Center |
| 56 | 10 SR-RIBKA = 235921 | Sweet/Grain Sorghum | Tigray |

**Supplementary Table 7.** Ablation study for training and loss function.

| Variant | In-distribution Test mIoU<br>(mean ± s.d., n = 185) | Out-of-distribution Test mIoU<br>(mean ± s.d., n = 35) |
| --- | --- | --- |
| *Encoder training strategy* |  |  |
| Frozen encoder | 0.881 ± 0.066 | 0.881 ± 0.034 |
| Fine-tuned encoder (RADIX) | 0.889 ± 0.065 | 0.888 ± 0.036 |
| *Loss function* |  |  |
| Dice + Focal + weighted CE | 0.885 ± 0.064 | 0.879 ± 0.038 |
| Dice + Focal + weighted CE + Lovasz (RADIX) | 0.889 ± 0.065 | 0.888 ± 0.036 |

## Supplementary Data

**Supplementary Data 1.**
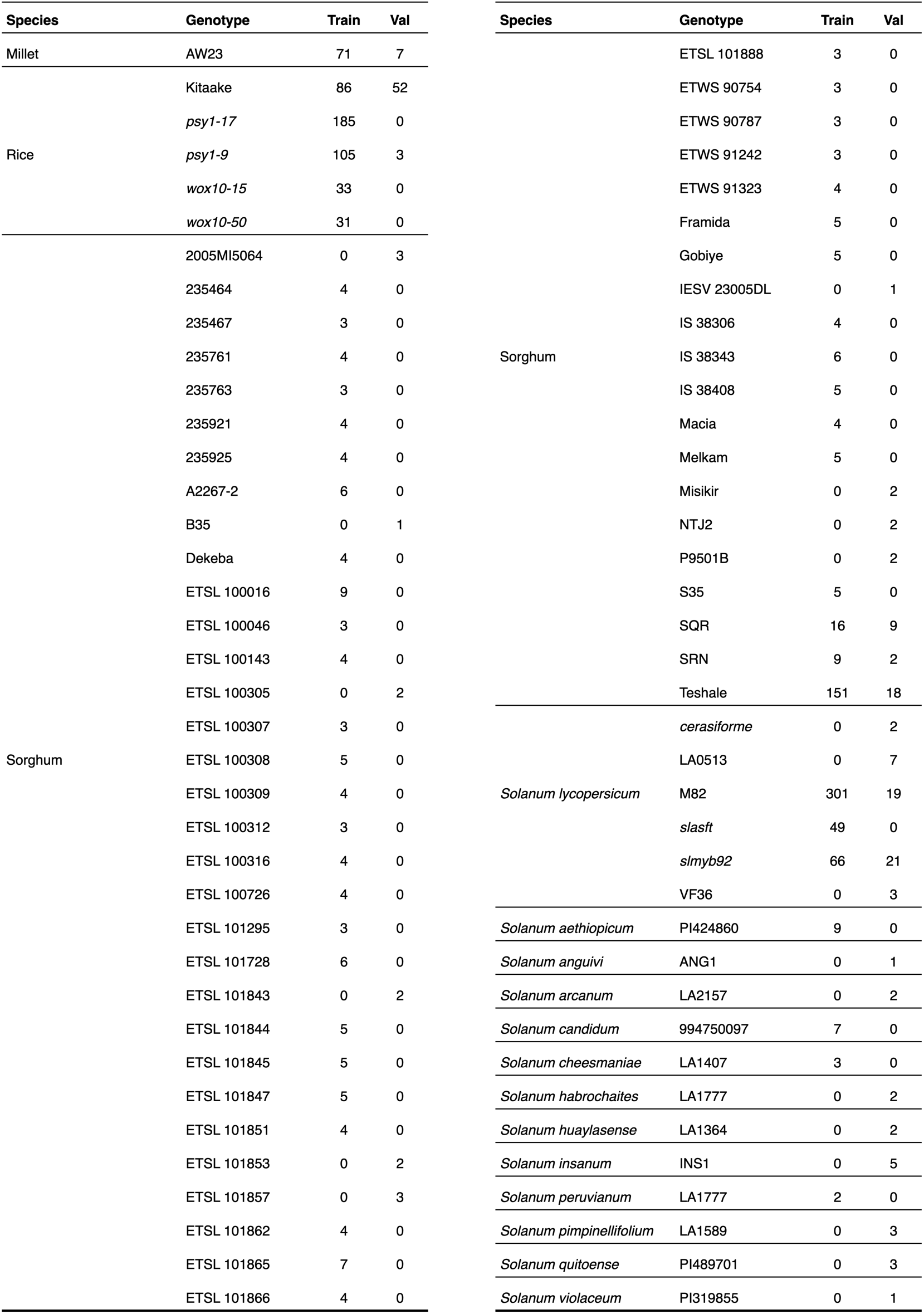
Training and validation set diversity (n = 1293 train, 182 val).

Supplementary Data 2. Imaging metadata

[Attached as an .xlsx file]

Supplementary Data 3. Annotation SOP

[Attached as an .docx file]

## SOP: Fluorescence Microscopy Image Annotation for Plant Root Segmentation

Authors: Stefan Sanow, Yifei Gu, Tamera Taylor, Kevin Morimoto

### 1. Purpose

This SOP provides standardized guidelines for generating consistent and accurate segmentation masks for tissue regions in fluorescence microscopy images of plant root cross-sections. These annotations are used to train and validate machine learning models that automatically identify and quantify key root structures. Annotations must be precise, as structural areas are measured in pixels.

### 2. Root Cross-Section Anatomy

#### Common Structures (All Species)

**Epidermis:** The outermost layer of the root, forming the interface between root tissue and the surrounding background. It appears as a layer of connected cells encircling the entire cross-section. The epidermis boundary defines the outer limit of the ’whole root’ annotation. In dicots (tomato), root hair extensions may be present and should be excluded from annotations.

**Exodermis:** The second outermost cell layer, lying directly beneath the epidermis and encasing the cortex. Exodermal cells are larger and more polygonal than epidermal cells. In multichannel images, the exodermis typically shows higher signal in two of three channels. It is annotated as two concentric rings: the outer exodermis (epidermis-facing boundary) and inner exodermis (cortex-facing boundary).

**Endodermis:** A thin layer of cells separating the cortex from the vascular cylinder. In dicots, it is the second full concentric layer from the vascular center (pericycle is first) and can be identified by distinct dot structures and banding patterns in multichannel images. It is annotated as two concentric rings: the outer endodermis (cortex-facing boundary) and inner endodermis (pericycle-facing boundary).

#### Monocot-Specific Structure (Millet, Rice, Sorghum)

**Aerenchyma:** Air-filled cavities that form within the cortex, appearing as dark, irregular holes between cells. They vary in size and shape. Not all monocot root images contain aerenchyma; roots without aerenchyma are annotated with zero polygons for label 1.

### 3. Annotation Classes

All species share the same 6-class label schema (indices 0–5). Classes that are biologically absent in a given species are annotated with zero polygons; this is real biology, not missing data.

| Label Index | Label | Description | Approximate Location / Notes |
| --- | --- | --- | --- |
| 0 | Whole Root | Outermost boundary of the root, separating sample from background. | Trace along the epidermis (outermost visible cell wall), ensuring the outermost cell wall is contained within the annotation mask. All species. |
| 1 | *Aerenchyma* | *Air spaces or cavities within the cortex.* | *Appear as dark, irregular holes in the cortex. Monocots (millet, rice, sorghum) only; annotate zero polygons in dicots.* |
| 2 | Outer Endodermis | Outer edge of the endodermis. | Trace the boundary between cortex and endodermis; contain the endodermis cell wall within the annotation mask. All species. |
| 3 | Inner Endodermis | Inner edge of the endodermis. | Trace the boundary between endodermis and pericycle; exclude the endodermis cell wall from the masked area. All species. |
| 4 | Outer Exodermis | Outer edge of the exodermis. | Trace the boundary between epidermis and exodermis; contain the exodermis cell wall within the annotation mask. All species. |
| 5 | Inner Exodermis | Inner edge of the exodermis. | Trace the boundary between exodermis and cortex; exclude the exodermis cell wall from the annotation mask. All species. |

Shaded row: biologically absent in dicots (zero polygons expected).

### 4. Annotation Guidelines

A. **General Rules**

1. Always follow visible cell walls when outlining boundaries.
2. Keep polygons clean, smooth, and non-overlapping.
3. When zooming in to annotate, periodically zoom out to ensure consistency across the full root section.
B. **Whole Root Boundary**

1. This is the outermost contour of the root cross-section.
2. Follow the epidermal layer tightly; do not include the background.
3. Ignore detached debris, root hairs (dicots), or broken edges outside the main root structure.
4. If a lateral root disrupts the boundary, draw across it to ensure a continuous closed polygon.
C. **Endodermis**

1. The outer and inner endodermis boundaries form concentric rings enclosing a narrow band of specialized cells.
2. **Outer endodermis (label 2):** annotate at the pixels just toward the cortex-facing side of the endodermal cells.
3. **Inner endodermis (label 3):** annotate at the pixels just toward the pericycle-facing side of the endodermal cells.
4. Maintain continuous, ring-like boundaries. If the line is dim, infer its likely continuation to complete the ring smoothly.
5. If a lateral root disrupts the endodermis, draw across it to keep both rings continuous.
D. **Exodermis**

1. The outer and inner exodermis boundaries form concentric rings enclosing the layer of cells directly beneath the epidermis.
2. **Outer exodermis (label 4):** annotate at the pixels just toward the epidermis-facing side of the exodermal cells.
3. **Inner exodermis (label 5):** annotate at the pixels just toward the cortex-facing side of the exodermal cells.
4. Maintain continuous, ring-like boundaries. If a line is dim, infer its likely continuation to complete the ring smoothly.
5. If a lateral root disrupts the exodermis, draw across it to keep both rings continuous.
E. **Aerenchyma - Monocots only (label 1)**

1. **Aerenchyma only occurs in the cortex.** Do not annotate gaps in the endodermis or epidermis — these are likely mechanical damage.
2. Annotate each distinct cortical cavity as one polygon, placed just inside the cellular boundaries.
3. Annotate small cavities individually if their shape is clearly defined.
4. A single polygon should not cover more than 25% of the total cortex area, unless the aerenchyma is very large and continuous.
5. If unsure whether a cavity is aerenchyma, annotate it — it is easier to remove uncertain annotations than to add missing ones.
6. Some images contain no aerenchyma; leave label 1 with zero polygons for those images.

### 5. Quality Control Checklist

Before finalizing annotations, verify that:

- All 6 label types are present and correctly indexed (0–5); biologically absent classes have zero polygons.
- All boundaries are smooth, closed polygons.
- No overlapping regions exist between classes.
- Aerenchyma annotations (label 1) are confined to the cortex in monocots; zero polygons in dicots.
- The root edge (label 0) follows the true epidermal boundary.
- Regions with ambiguous gaps or broken cells have been double-checked.

### 6. Annotation Format

Export annotations in **YOLO-style segmentation format** as .txt files. Each line contains the label index followed by normalized x,y coordinate pairs defining the polygon vertices:

<label_index> x1 y1 x2 y2 x3 y3 …

#### Example (label 2 — Outer Endodermis)

2 0.399595 0.479795 0.402324 0.477905 0.405264 0.476646 0.407783 0.474546 …

## Notes

### Competing Interest Statement

The authors have declared no competing interest.

